# Secreted inhibitors drive the loss of regeneration competence in *Xenopus* limbs

**DOI:** 10.1101/2020.06.01.127654

**Authors:** C. Aztekin, T. W. Hiscock, J. B. Gurdon, J. Jullien, J. C. Marioni, B. D. Simons

## Abstract

Absence of a specialised wound epidermis is hypothesised to block limb regeneration in higher vertebrates. To elucidate the cellular and molecular determinants of this tissue, we performed single-cell transcriptomics in regeneration-competent, -restricted, and -incompetent *Xenopus* tadpoles. We identified apical-ectodermal-ridge (AER) cells as the specialised wound epidermis, and found that their abundance on the amputation plane correlates with regeneration potential and injury-induced mesenchymal plasticity. By using *ex vivo* regenerating limb cultures, we demonstrate that extrinsic cues produced during limb development block AER cell formation. We identify *Noggin*, a morphogen expressed in cartilage/bone progenitor cells, as one of the key inhibitors of AER cell formation in regeneration-incompetent tadpoles. Extrinsic inhibitory cues can be overridden by *Fgf10*, which operates upstream of *Noggin* and blocks chondrogenesis. Together, these results indicate that manipulation of the extracellular environment and/or chondrogenesis may provide a strategy to restore regeneration potential in higher vertebrates.

**One Sentence Summary:** Extrinsic cues associated with chondrogenic progression inhibit AER cell formation and restrict limb regeneration potential.

## Introduction

Amphibian limb regeneration relies on a specialised wound epidermis (also known as the apical-epithelial-cap, AEC). It has been hypothesised that the absence of this tissue limits the regeneration potential of higher vertebrates, including mammals (*1*). The AEC was suggested to be analogous to the apical-ectodermal-ridge (AER) that is specifically seen during limb development, since both tissues are required for proximal-distal outgrowth, and express *Fgf8* at the distal tips of limb buds/amputation planes (*2*). It has been proposed that the AER functions to maintain and enable proliferation of underlying cells in distal mesoderm. Similarly, the AEC was hypothesised to be formed upon amputation in order to enable the self-renewal of underlying progenitor and dedifferentiated cells, leading to the formation of a proliferative structure called the blastema (*3*). Injury was thought to induce dedifferentiation of cells that can interact with AEC to build a blastema (*4*), although it is not clear how a specialised wound epidermis forms during regeneration and why it cannot form in some instances/species.

*Xenopus laevis* tadpoles lose limb regeneration ability progressively during their development, coinciding with their inability to form a specialised wound epidermis (*5, 6*). At the developmental stages prior to the formation of digits, amputations lead to a complete regeneration of the limb (Niewkoop & Faber (*7*) (NF) ∼52-54, regeneration-competent). As autopod development proceeds, amputations result in partial regeneration, characterized by missing digits (NF ∼55-57, regeneration-restricted). Towards metamorphosis, amputations either cause the growth of a spike-like cartilaginous structure without joints and muscles, or a simple wound healing (NF ∼58 and beyond, regeneration-incompetent). Moreover, *Xenopus* limb regeneration ability declines when amputations are performed at more proximal regions of the limb, where there are more mature chondrogenic and osteogenic cells (*8, 9*). Likewise amputation through bone results in reduced regeneration compared to amputations at the joints (*8, 9*). Nevertheless, certain procedures (*e.g.* FGF8 or FGF10 bead applications) can induce specialised wound epidermis formation and restore, or contribute to, limb regeneration in otherwise regeneration–incompetent species (*10, 11*). Therefore, investigating the cellular and molecular mechanisms controlling specialised wound epidermis formation can help to devise new strategies to promote mammalian limb regeneration.

### Single-cell RNA-seq analysis reveals cell type heterogeneity during development and following amputation of the limb

To characterise cellular changes associated with regeneration-ability, we first sequenced developing intact hindlimbs at particular morphologically-defined stages: NF Stage ∼52 (limb bud stages), NF Stage ∼54 (autopod forming) and NF Stage ∼56 (autopod formed) **(Fig. 1A).** Then, to evaluate the cellular responses to injury and regeneration, we profiled cells from amputated limbs and their contralateral controls. Specifically, we amputated hindlimbs from presumptive knee/ankle levels for regeneration-competent tadpoles (NF Stage ∼52-53) and ankle level for –restricted (NF Stage ∼55-56) and –incompetent tadpoles (NF Stage ∼58-60), and sequenced cells from newly-generated tissues at 5 days post-amputation (dpa) **(Fig. 1B)** when the specialised wound epidermis and blastema are seen morphologically (*2*). Contralateral developing limb buds or autopods were sequenced as controls. We did not include a contralateral control at the regeneration-incompetent stage as we were not able to obtain cells with the dissociation protocol used to process the other samples.

**Fig 1.**
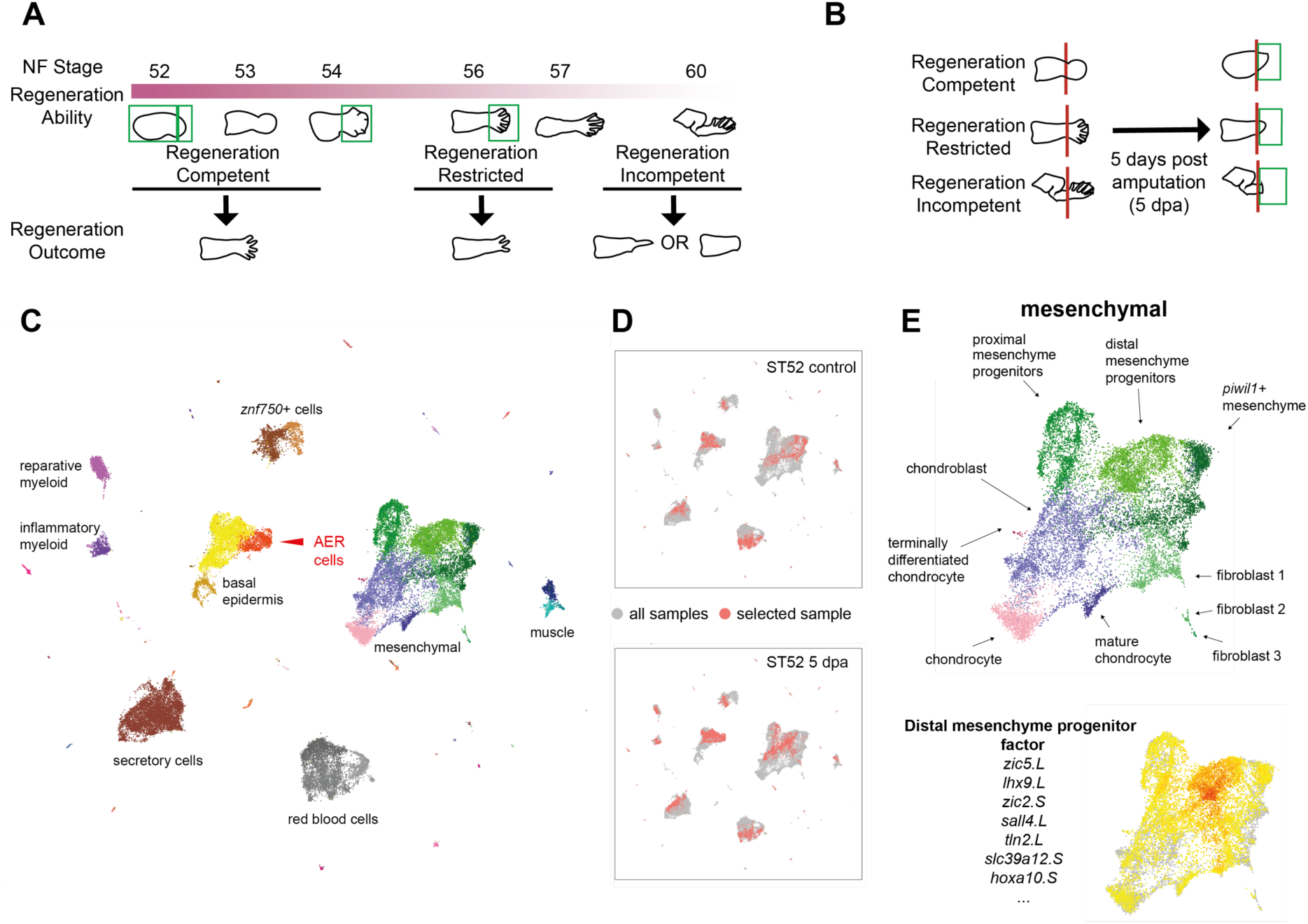
Single-cell transcriptomics reveals cellular heterogeneity in developing and amputated *Xenopus* limbs at different stages of regeneration competence. **A)** Schematic describing *Xenopus* limb regeneration at different NF Stages. NF Stage ∼52-54 tadpoles are regeneration-competent and amputations result in regeneration of a full limb. Regeneration-ability begins to decline at NF Stage ∼54. Tadpoles are regeneration-restricted at NF ∼Stage 56 where 2-3 digits can be regenerated. Beyond NF Stage ∼58, tadpoles are regeneration-incompetent and amputations result in simple wound healing or unpatterned spike formation. Green boxes indicate the samples collected for scRNA-Seq, taken at stages prior to, at the onset of and after the loss of regeneration ability. **B)** Schematic describing 5 days post amputation (dpa) samples for regeneration-competent, -restricted, and –incompetent tadpoles. Green boxes show the samples collected for scRNA-Seq. **C)** An atlas of cell types in intact and amputated limbs. Samples from each condition are processed separately for sequencing, and are then pooled together for UMAP visualization and clustering. Each dot corresponds to a single cell, colours indicate cluster identity, text labels highlight important tissue/cell types. See Fig. S3 for full annotation. **D)** Comparisons can be made between conditions to highlight transcriptional changes associated with regeneration; here NF Stage 52 amputated limbs (lower) are compared to their contralateral control samples (upper). Red dots denote cells in the selected sample; grey dots denote cells in all samples. **E)** A diversity of mesenchymal cell types is detected in our dataset (upper), together with putative gene expression programs identified using unbiased factor analysis (lower).

Next, we pooled the single-cell RNA sequencing data derived from at least two replicates for each condition **(Fig. S1)**, corrected our atlas for cell cycle effects **(Fig. S2)**, resulting in a total of 42,348 cells **(Methods; Fig. 1C-D, Fig. S3-4**). Following clustering of cells based upon their gene expression profiles, examination of multiple marker genes **(Fig. S5)** revealed at least 60 distinct clusters representative of putative cell types **(Fig. 1C and Fig. S3)** including known populations (e.g. AER cells) and potentially new uncharacterised cell states (e.g. a *Piwil1*+ population in the mesenchyme) **(Fig. 1E).** From the cell atlas, we were able to detect cell cycle differences between cell types, e.g. distal mesenchyme progenitors were more biased towards G2/M phases compared to proximal mesenchyme progenitors **(Fig. S2D)**, as reported in mouse (*12*). The *Xenopus* limb cell atlas is accessible using an interactive platform (https://marionilab.cruk.cam.ac.uk/XenopusLimbRegeneration/).

### Quantitative features of AER cell formation are associated with regeneration outcome

We then focused on the specialised wound epidermis, or AEC, that was suggested to be analogous to the AER. Although both populations were characterized by *Fgf8* expression (*2*), the extent of similarity between these cells was unclear. In our data, we could detect mostly quantitative gene expression differences between cells defined as belonging to the AER (defined as *Fgf8* expressing epidermal cells during limb development) and the AEC (*Fgf8* expressing epidermal cells in 5 dpa samples) (**Fig. 2A-B).** Moreover, cells related to these tissues were aggregated in the same *Fgf8*+ epidermal cluster (**Fig. 2B-C**). Additionally, both during development and post-amputation, 5 dpa *Fgf8+* epidermal cells were mostly detected as a monolayer of polarised cuboidal basal cells, **(Fig. S6**) though multilayers were seen to form in some instances (**Fig. S7)**. Hence, based on their transcriptomic signature, tissue localisation, and general cellular morphology, the cells populating the AER structure during development and the AEC specialised wound epidermis structure found during regeneration were referred to as *AER cells* in this study. Finally, we found that AER cells (limb specialised wound epidermal cells) and cells that define the specialised wound epidermis during *Xenopus* tail regeneration (regeneration-organising-cells, ROCs (*13*)) showed similar, but non-identical gene expression profiles **(Fig. S8)**, emphasizing that different cell types operate in different appendage regeneration scenarios.

**Fig 2:**
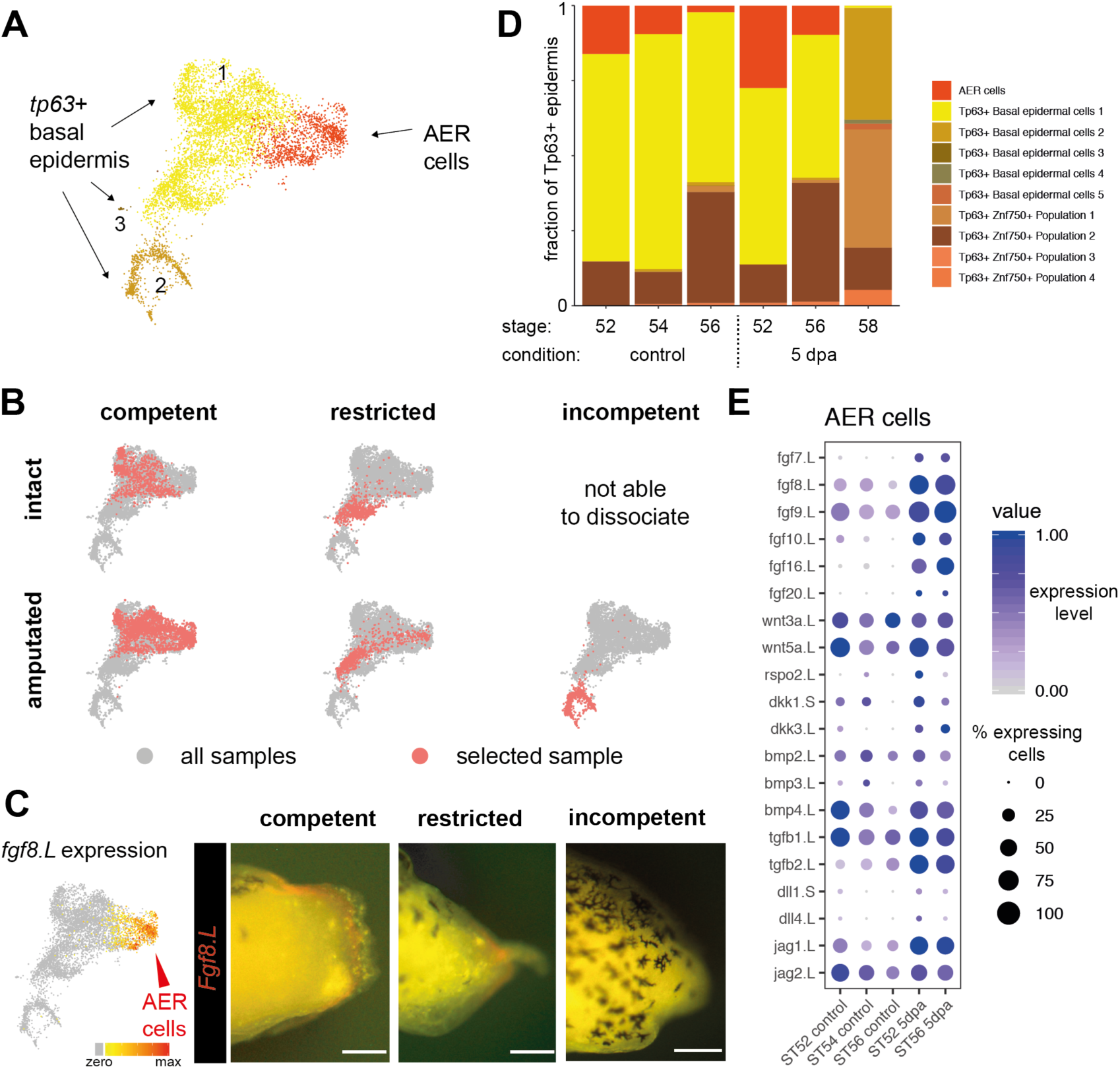
Formation of a signalling centre comprising apical-ectodermal-ridge (AER) cells is associated with the successful regeneration. **A)** Multiple basal epidermal cell states are detected, including AER cells. **B)** UMAP visualisation of basal epidermis reveals that re-establishment of AER cells is associated with successful regeneration. Red dots denote cells in the selected sample; grey dots denote cells in all samples. **C)** (Left) AER cells express *Fgf8.L*. (Right) Stereomicroscope images of the 5 dpa amputation plane of regeneration-competent, restricted, and –incompetent tadpoles. *Fgf8.L* (red) expressing AER cells are formed in regeneration-competent and –restricted tadpoles, but not in –incompetent tadpoles. Scale bar = 250 µm. **D)** Abundance of basal epidermal cell types across conditions reveals a correlation between AER abundance and regeneration outcome. AER cells are present in intact –competent samples, and are enriched after amputation. A similar pattern is seen in -restricted samples, although abundances of AER cells are reduced. Very few AER cells are detected in –incompetent tadpoles. **E)** Dot plot showing expression of selected ligands for AER cells during development and at 5 dpa regeneration-competent, restricted, and –incompetent samples. Dot colour indicates mean expression; dot size represents the percentage of cells with non-zero expression.

Limb amputation results in the formation of *Fgf8* expressing AEC at the amputation plane in regeneration–competent tadpoles, but not in –incompetent tadpoles (*6*), while AEC formation has not been characterised previously for –restricted tadpoles. Using our atlas, we found that, at 5 dpa, tadpole epidermis contained abundant AER cells in regeneration–competent tadpoles, a limited number of AER cells in -restricted tadpoles, and that AER cells were largely absent from -incompetent tadpoles **(Fig. 2B-D).** The signalling centre properties of AER cells was reflected in the many diverse ligands they express, which can influence proliferation and cell fate decisions **(Fig. 2E and Fig. S9).** Although *Fgf8* was always expressed in AER cells, relative expression of *Fgf8* and other ligands varied among conditions **(Fig. 2E)**. Overall, while the signalling centre potency of AER cells appears variable, a strong correlation between AER cell abundance and regeneration-outcome was evident.

### The presence of AER cells has an association with injury-induced mesenchymal plasticity

It has been suggested that the AEC enables the self-renewal activity of dedifferentiated cells, leading to blastema formation (*3*). To identify signatures of dedifferentiation in our atlas, we first examined the expression of genes related to dedifferentiation and blastema formation (e.g. *Sall4, Kazald1, Marcksl1* (*14*)). We observed that these genes were found to be either already expressed before amputation or upregulated upon amputation in a subset of fibroblasts **(Fig. S10A-B)**, which were located near the skin and perichondrium **(Fig. S11)**. Likewise, we found that a small fraction of these fibroblasts expressed muscle-related genes (e.g. *Pax3*) before and after amputation **(Fig. S10B)**. Moreover, independent of regeneration-outcome, amputation resulted in these fibroblast cells to express genes related to distal mesenchyme progenitors (e.g. *Grem1, Shh, Msx1, Fgf10*), and chondrogenesis (e.g. *Col8a2, Sox9)* **(Fig. S10A)**. Lastly, amputation not only increased the expression of known marker genes, but also up-regulated the expression of an entire putative distal mesenchyme progenitor gene set **(Fig. S10C)**, with the magnitude of this expression being lower in samples having fewer AER cells. Together, we concluded that, upon amputation, a subset of fibroblasts manifest injury-induced mesenchymal plasticity, the extent of which tracks with AER cell abundance.

### AER cell formation requires activation of multiple signalling pathways

To investigate the molecular mechanisms that mediate AER cell formation upon amputation, we developed an *ex vivo* regenerating limb culture protocol, inspired by previous work (*15*) **(Fig. 3A)**. By culturing amputated stylopod or zeugopod/stylopod from regeneration-competent and –restricted tadpoles, respectively, we observed *Fgf8* cell formation at the distal part of explants within 3 dpa (**Fig. 3B**). These explants also exhibited cone-shaped growth as cells accumulated uniformly underneath *Fgf8* cells, mimicking *in vivo* regeneration **(Fig. 3A-B, Fig S12B).** Interestingly, the proximal site of explants was also covered with epidermis **(Fig. S13A)**, but neither *Fgf8* expressing cells nor a uniform cell accumulation underneath the epidermis was observed **(Fig. 3A-B, S12B).** Moreover, the proximal site of the explant exhibited active chondrogenesis, manifesting in an outwards growth of cartilaginous tissue **(Fig. 3A, Fig S12C)**. This phenotype was particularly pronounced when explants were harvested from developmental stages in which proximal tissues were advanced in chondrogenesis (onset of NF Stage 53-54) **(Fig. S12D)**, and could be further enhanced by addition of BMP4, a known chondrogenesis inducer **(Fig S12E)**. Hence, the proximal and distal sites of limb explants exhibit different behaviours: the distal sites recapitulate localised AER cell formation as seen *in vivo*, while the proximal site is characterised by active chondrogenesis.

**Fig 3.**
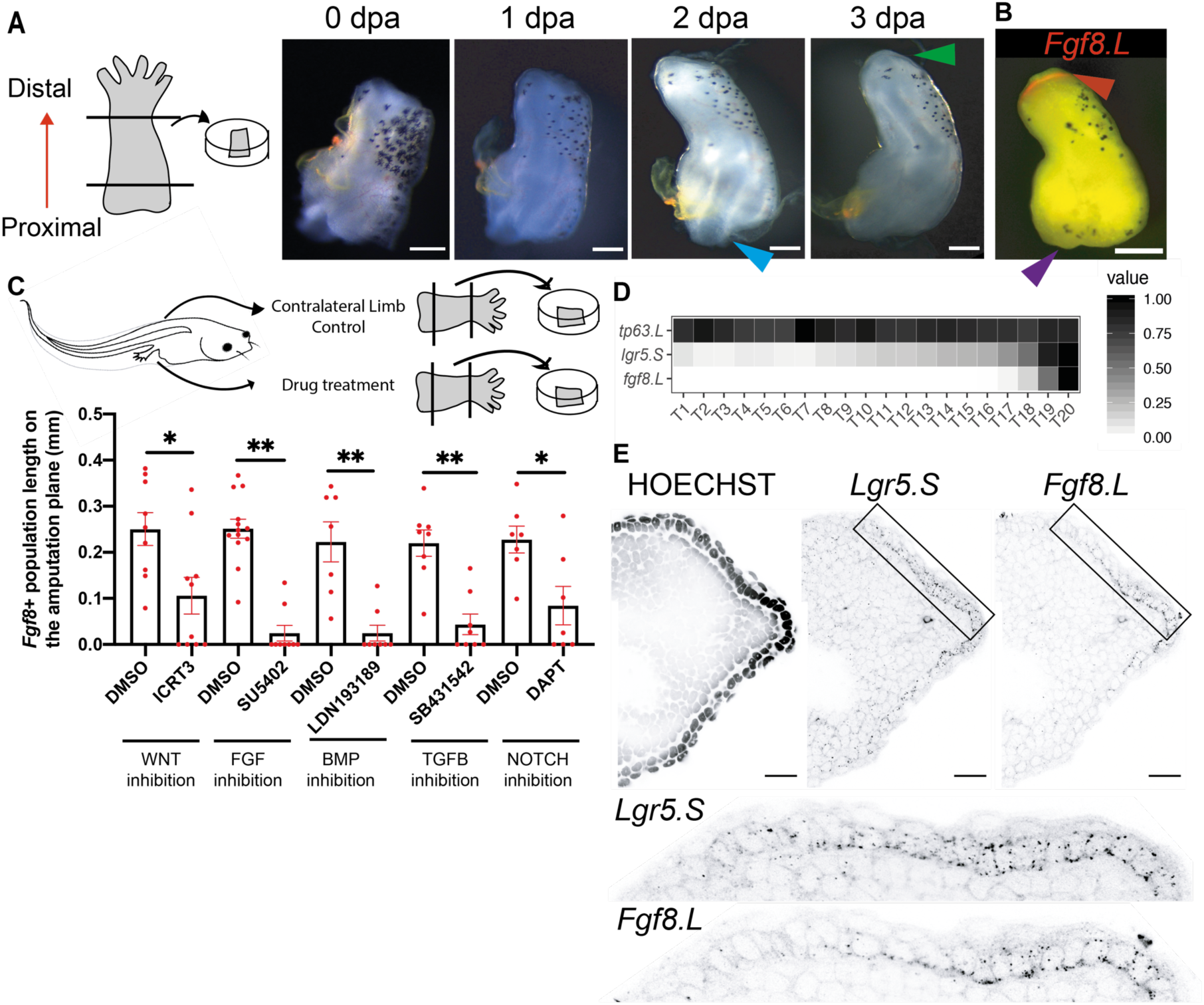
*Ex vivo* regenerating limbs demonstrate that AER cell formation requires activation of multiple pathways and can form from basal epidermal cells. **A)** (Left) Schematic for *ex vivo* regeneration limb culture. (Right) Time-lapse images of a – competent explant. The explant grows a cone shape at its distal site reminiscent of *in vivo* regeneration (green arrow), whilst the proximal site shows chondrogenesis (blue arrow). Scale= 200 µm. **B)** Example image of a –competent explant at 3-day post culture. Distal site of explants is *Fgf8.L* positive (red arrow), and proximal site is *Fgf8.L* negative (purple arrow). Red, *Fgf8.L* mRNA. Scale= 200 µm. **C)** Drug screen to test regulators of AER cell formation. (Top) Schematics describing the screen. One limb of a tadpole was used for perturbation and the contralateral limb from the same tadpole was used as a control. Samples were treated with the indicated drugs for 3 days post culture, and then stained for *Fgf8.L* mRNA. The extent of *Fgf8.L* expression along the amputation plane was measured. Sample sizes: ICRT3 total n≥9 from 3 biological replicates; SU5402 total n≥9 from 2 biological replicates; LDN193189 total n=8 from 3 biological replicates; DAPT total n=7 from 3 biological replicates. *P*\*< 0.05, and *P*\*\*< 0.001. **D)** Factor analysis identifies a putative gene expression trajectory from basal epidermal cells to AER cells predicting sequential activation of *Lgr5.S* followed by *Fgf8.L*. **E)** A proximal-to-distal gradient of *Lgr5.*S and *Fgf8.L* is observed in *vivo*, with *Fgf8.L* being restricted to the most distal regions of the midline epidermis. Black dots represent HCR mRNA signal. Scale = 20 µm.

In addition to changes associated with regeneration, explants could be used to determine signalling requirements for specialised wound epidermis formation. Inhibition of FGF, BMP, and WNT pathways via small molecule inhibitors blocked AER cell formation in explants **(Fig. 3C)**, reinforcing the conclusion that the reported *in vivo* AEC effects are mediated through a direct effect on the limb rather than a systemic effect (*16–18*). Moreover, by using the culture assay, we found that active TGF-β and NOTCH signalling are also required for *Xenopus* AER cell formation **(Fig. 3C)**. Overall, we concluded that AER cell formation requires the activity of multiple major signalling pathways.

### AER cells can form without cell division

Having established the molecular pathways required for AER cell formation, we asked how AER cells form on the amputation plane. By tracing skin tissue located on the edge of explants we found that they contributed to the covering of both the distal and proximal sites **(Fig. S13B)**. As the amputation planes are covered by skin tissue from the surrounding area, we reasoned that AER cells are likely to have originated from skin cells. As amputation eliminates the majority, if not all, of AER cells in the limb, we hypothesized that AER cells are derived from remaining skin stem cells. If AER cells are induced through proliferation and differentiation following amputation, all AER cells should be the product of cell division. To test this, we assayed the level of EdU incorporation in newly-formed AER cells. Surprisingly, we found that only ∼40% of AER cells (distal epidermal *Fgf8*+) were EdU positive at 3 dpa **(Fig. S13C)**, suggesting that most AER cells are induced independent of cell division following amputation. Consistently, a step-wise activation of *Lgr5.S* (a WNT target gene) followed by *Fgf8.L* expressions was identified as a possible gene-expression trajectory that could allow basal epidermal cells to convert directly to AER cells (**Fig. 3D).** Consistent with such a process, when visualized *in vivo*, we found that *Fgf8*+*/Lgr5*+ AER cells were flanked by *Lgr5*+ cells in the basal epidermis on the amputation plane or in the developing limb **(Fig. 3E, Fig. S6)**. Overall, these results support the hypothesis that basal epidermal cells can acquire AER cell identity without cell division.

### Loss of regeneration potential is associated with extrinsic cues inhibitory to AER formation

We then asked why fewer, or no, AER cells form on the amputation plane of regeneration-restricted or -incompetent tadpoles, respectively. Previous studies have shown that addition of *Fgf10* can induce AER formation in otherwise regeneration-incompetent tadpoles (*10*), suggesting that epidermal cells in these animals are intrinsically competent to form AER cells. Hence, we focused on the possibility that extrinsic cues are the basis for a reduced regeneration potential.

To test whether environmental factors secreted from -incompetent tadpoles would block AER cell formation, we took advantage of our *ex vivo* cultures. First, we co-cultured *ex vivo* limbs from regeneration-competent and –incompetent tadpoles. Strikingly, when such cultures were stained against *Fgf8* at 3 dpa, we observed that regeneration-competent tadpole limbs failed to form AER cells **(Fig. 4A**). Second, we collected media from regeneration-incompetent tadpole explants and cultured freshly amputated regeneration-competent explants with this conditioned media. Consistent with the co-culture experiment, the conditioned media from regeneration-incompetent tadpoles blocked AER cell formation in -competent explants **(Fig. 4B)**. By contrast, neither co-culturing with regeneration–competent explants, nor preparing conditioned media from regeneration-competent explants, affected AER cell formation in regeneration– competent explants **(Fig 4A-B)**. Additionally, conditioned media from regeneration– competent explants was unable to induce AER cell formation in –incompetent explants **(Fig S14)**. Altogether, these results suggest that secreted inhibitory factors block AER cell formation in regeneration-incompetent tadpoles compromising their regeneration potential.

**Fig 4.**
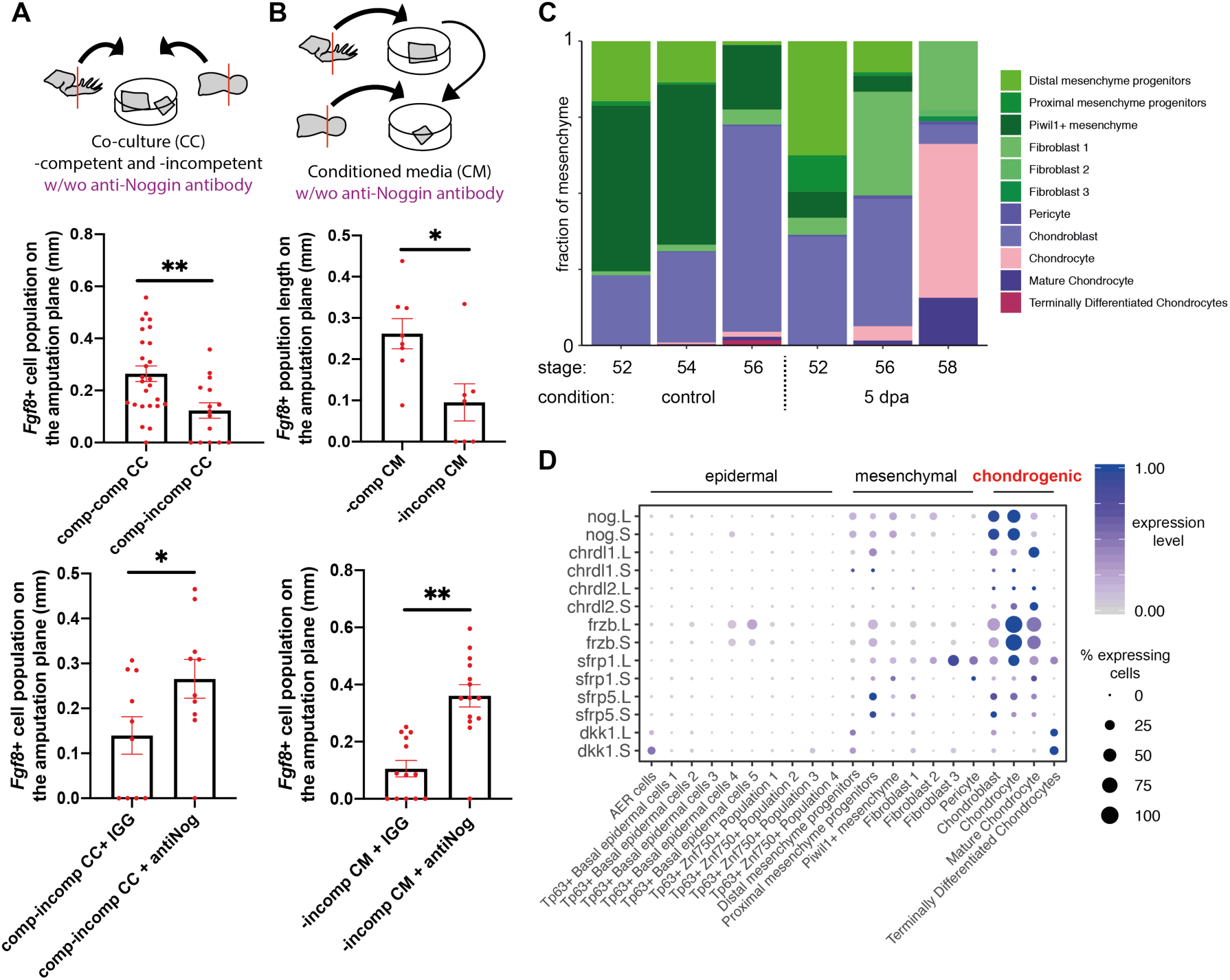
Inhibitory factors, such as Noggin, are secreted from chondrogenic populations at regeneration incompetent stages, and block AER cell formation. **A)** (Top) Schematic describing conditioned media experiments to test the effect of extrinsic cues in regeneration-incompetent tadpole limbs. (Mid) Supplying conditioned media (CM) from regeneration-incompetent tadpoles to regeneration-competent explants decreases the extent of *Fgf8.*L expression at the amputation plane at 3 dpa. (Bottom) This effect can be rescued by adding anti-NOGGIN antibody. -Competent CM to –competent explants: total n=8, from 3 biological replicates; -incompetent CM to –competent explants: total n=7, from 3 biological replicates; -incompetent CM to –competent explants and anti-IGG antibody: total n=10, from 3 biological replicates; -incompetent CM and anti-NOGGIN antibody to – competent explants: total n=10, from 3 biological replicates. *P*\*< 0.05, and *P*\*\*< 0.001. **B)** (Top) Schematic describing co-culture experiments. (Mid) Co-culturing (CC) -competent explants and –incompetent explants decrease the extent of *Fgf8.*L expression at the amputation plane at 3 dpa. (Bottom) This effect can be rescued by adding anti-NOGGIN antibody. – Competent and –competent CC: total n=26, from 4 biological replicates; -competent and – incompetent CC: and anti-IGG antibody total n=10, from 3 biological replicates; competent and –incompetent CC and anti-NOGGIN antibody: total n=10, from 3 biological replicates. *P*\*< 0.05, and *P*\*\*< 0.001. **C)** Abundance of mesenchymal populations across conditions reveals an enrichment of chondrogenic populations at regeneration-restricted and -incompetent stages, in both intact and amputated limbs. **D)** Multiple BMP/WNT antagonists are expressed specifically in chondrogenic populations.

To identify the factors responsible for this inhibitory effect, we surveyed our single-cell atlas for the expression of secreted proteins involved in signalling pathways required for AER cell formation. We found that that the loss of regeneration potential is associated with an increased proportion of chondrogenic lineage cells in the mesenchyme **(Fig. 4C)**, and that these cells express multiple inhibitory ligands for BMP and WNT pathways **(Fig. 4D).** As chondrogenic populations specifically express high levels of *Noggin* **(Fig. 4D)**, a known antagonist of BMP signalling, we hypothesised that AER cell formation is antagonised by an excess of secreted *Noggin* in regeneration-incompetent tadpoles. Indeed, consistent with previous observations (*19*), addition of NOGGIN to regeneration-competent *ex vivo* limbs blocked AER cell formation **(Fig. S15A).** To test if *Noggin* does indeed act as one of the inhibitory extrinsic cues produced following amputation in regeneration-incompetent tadpoles, we blocked NOGGIN in our co-culture and conditioned media experiments using anti-NOGGIN antibodies **(Fig. 4A-B)**. Strikingly, blocking secreted NOGGIN by antibody addition cancelled the inhibitor*y* activity on AER cell formation in both co-culture and conditioned media experiments **(Fig. 4 A-B)**.

As these experiments point towards the chondrogenic lineage as the source of inhibitory extrinsic cues, we then asked if limiting chondrogenesis can promote AER cell formation. To this end, we generated tip explants by culturing distal limb buds (NF Stage ∼52) or early formed autopods (NF Stage ∼54) without their proximal segment, where the most advanced chondrogenesis takes place. Indeed, these tip explants showed ectopic *Fgf8* expression at different sites of the epidermis further suggesting a localised and/or long-range inhibitory effect of secreted factors from mature chondrogenic cells (**Fig. S15B)**. Overall, these results indicate that the loss of regeneration ability is associated with extrinsic cues, including NOGGIN, inhibitory to AER cell formation.

To test if manipulation of BMP signalling can also enhance AER cell formation in regeneration–competent tadpoles, we perturbed the BMP pathway. Indeed, we found that inhibiting NOGGIN could increase the formation of AER cells **(Fig. S15A**). By contrast, the addition of BMP4 to regeneration–competent *ex vivo* cultures blocked AER cell formation **(Fig. S15A)**, an effect similar to that reported in AER development in chick embryos (*20, 21*). As BMP4 boosts chondrogenesis **(Fig. S12)**, which can in turn lead to *Noggin* expression, we concluded that localised and/or regulated levels of BMP agonist and antagonists are key for AER cell formation.

### FGFR activation negatively regulates progression of chondrogenesis and FGF pathway operates upstream of Noggin for AER cell formation

Regeneration competency in late stage tadpoles was previously shown to be restored via exogenous application of FGF10, which activates the expression of genes associated with regeneration and AER cells (*10*) (**Fig. S16A)**. However, the mechanism by which FGF10 regulates the formation of AER cells is not clear. We first asked if all cells in the epidermis are competent to induce *Fgf8* expression upon *Fgf10* exposure. We examined the spatial correlation between *Fgf10.L* expressing mesenchymal cells and *Fgf8.L* expressing epithelial cells in regeneration–competent tadpoles and saw regions in which *Fgf10.L* but not *Fgf8.L* was present **(Fig. S16B)**. Second, when adding FGF10 to regeneration–competent explants, we observed a slight but not statistically significant increase in AER cell formation on the amputation plane **(Fig. S16C)**, although this signal was confined to the distal epidermis and did not include a substantial signal at the proximal site of explants **(Fig. S16D)**. Collectively, we concluded that *Fgf10* is not sufficient to induce AER cells across the whole epidermis. We next sought to evaluate if the effect of *Fgf10* on regeneration is, at least in part, mediated by its impact on chondrogenesis.

To test the effect of *Fgf10* on chondrogenesis, we used our *ex vivo* cultures to monitor the substantial chondrogenesis occurring at the proximal site of explants. Application of FGF10 beads to the proximal site of *ex vivo* cultures, or addition of recombinant FGF10 to their media, significantly decreased chondrogenesis at the posterior sites in regeneration-restricted explants **(Fig. S17A).** Conversely, blocking FGFR significantly extended chondrogenesis at the proximal site of explants **(Fig. S17B-D).** Nonetheless, FGF10 treatment was not sufficient to induce strong *Fgf8* expression at the proximal site of explants **(Fig. S16D)**, presumably due to abundant antagonist cues. To test this hypothesis, we treated explants with a combination of FGF10 and anti-NOGGIN antibodies. Strikingly, this combination not only enhanced AER cell formation at the distal sites, but also induced ectopic *Fgf8.L* expression near the proximal sites of explants **(Fig. S16D)**. Finally, AER cell formation induced by FGF10 addition was cancelled by the addition of BMP inhibitors (NOGGIN or small molecule inhibitors) **(Fig. S16C)**, further suggesting that FGF10 acts events upstream of potential inhibitory extrinsic cues during AER cell formation.

## Discussion

Here, we have used single-cell transcriptional profiling to discriminate injury responses from regeneration-specific events in *Xenopus* limbs. Consistent with previous observations based on tissue-level characterisations (*4*), using the single-cell atlas, we found that the abundance of AER cells on the amputation plane correlates with regeneration outcome and injury-induced mesenchymal plasticity.

Changes in intrinsic properties of mesodermal tissue during development, particularly the loss of *Fgf10* expression, was suggested to explain regeneration-incompetency (*22, 23*). Our results suggest that extrinsic cues from mesodermal tissue also affect regeneration strongly by inhibiting specialised wound epidermis formation, downstream of *Fgf10* signalling. Specifically, chondrogenic cells are the main source of secreted extrinsic cues (*e.g. Noggin*) that block AER cell formation. These findings may explain why amputations at proximal versus distal sites, associated with different stages of chondrogenesis, exhibit different regeneration outcomes (*9*).

Whilst manipulation of chondrogenesis in adult frogs and other regeneration–incompetent species may lead to novel approaches to promote limb regeneration, it is likely that additional barriers to regeneration (*e.g.* scarring and more complex immune responses) will also have to be overcome. Finally, it is tempting to speculate whether limb regeneration-competent salamanders can withstand the inhibitory extrinsic cues by having AER cell signals enriched in mesenchymal rather than epidermal cells, or whether they utilise different mechanisms owing to distinct limb development mechanisms (*24–26*). Altogether, our work suggests a new cellular model of limb regeneration **(Fig 5)**, which unites disparate findings in the field, and suggests that modulation of extrinsic cues impacting on epidermal populations has the potential to unlock the ability to regrow lost limbs in non-regenerative higher vertebrates.

**Fig 5.**
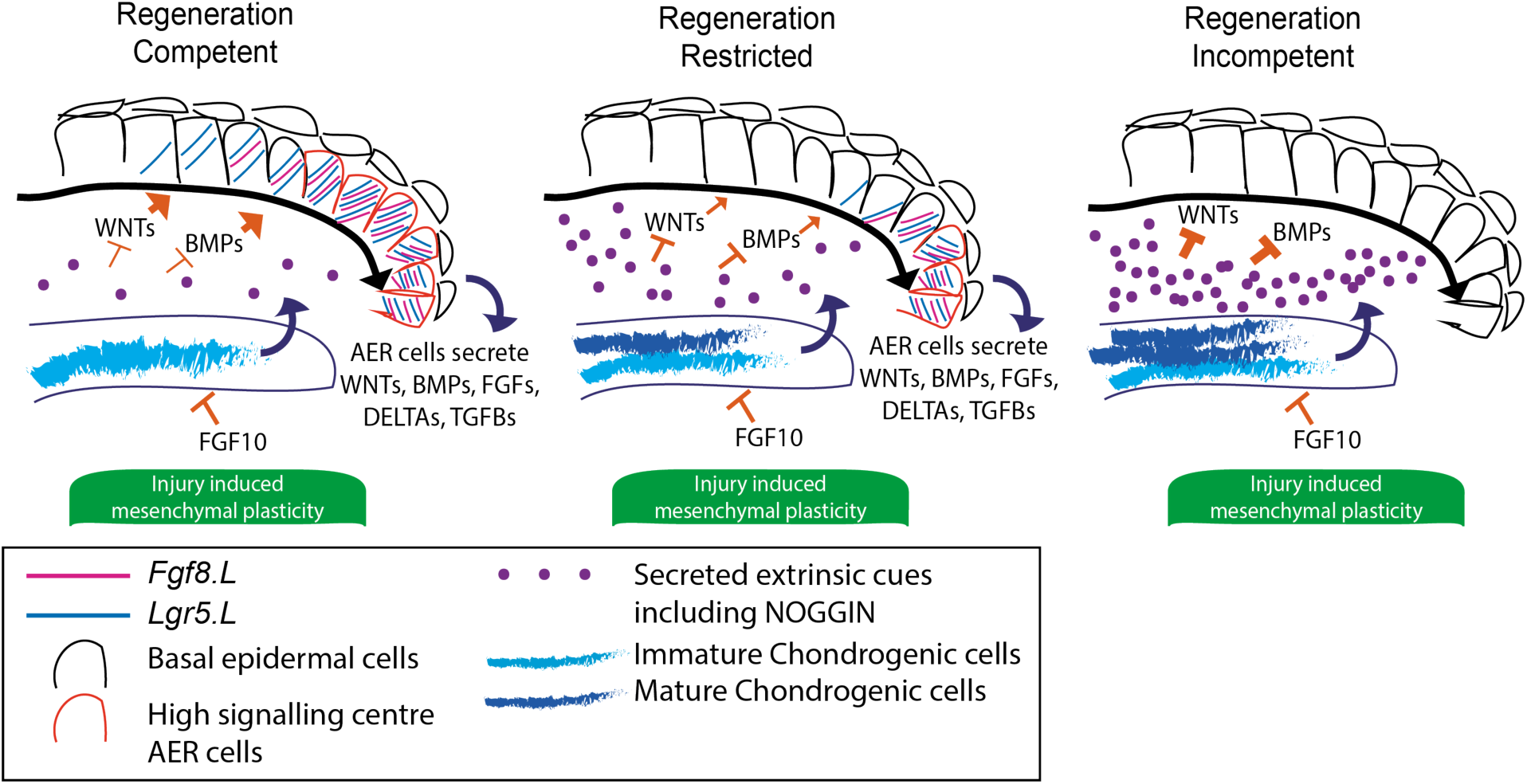
Chondrogenic progression blocks AER cell formation by modulating extrinsic cues. Secreted factors such as WNTs and BMPs support AER cell formation at the amputation plane. During development, chondrogenesis leads to the accumulation of secreted inhibitory extrinsic cues including NOGGIN which results in failure to establish AER cells (*Fgf8.L+/Lgr5.S+).* FGF10 can suppress chondrogenesis. Amputations, independent of the regeneration outcome, induce injury-induced mesenchymal transcriptional plasticity.

## Acknowledgements

We thank Katarzyna Kania and the Cambridge Institute Genomics Core for their support with this work on the 10X-Genomics and sequencing library preparations. The transgenic testes used in this study was obtained from the European Xenopus Resource Centre, curated with funding from the Wellcome Trust/BBSRC, and maintained by the University of Portsmouth, School of Biological Sciences. We thank R. Jones-Green for excellent animal care. We thank H. Ma for use of her stereomicroscope, and E. Rawlins for use of her Zeiss Axiolmager compound microscope. We thank B. Steventon, E. Rawlins, and members of the Marioni and Simons labs for general discussion about the project. We thank R. Butler for assistance in image analysis.

## Author contributions

Conceptualization: C.A. with contributions from other authors; Methodology: C.A., and T.W.H. for computational analysis; Software: T.W.H.; Validation: C.A., T.W.H.; Formal analysis: T.W.H. with help from C.A.; Investigation: C.A.; Resources: J.B.G., J.C.M.; Data curation: C.A., T.W.H; Writing – original draft: C.A. with help from J.J.; Writing – review and editing: C.A., T.W.H, J.J., B.D.S., J.C.M.; Supervision: J.C.M., J.J., B.D.S.; Project administration: C.A.; Funding acquisition: C.A., J.J., J.B.G., B.D.S., J.C.M.

## Data and materials availability

Sequencing data and processed gene counts are available on ArrayExpress with the accession number E-MTAB-9104. Analysis scripts are available at https://github.com/MarioniLab/XenopusLimbRegeneration2020. Requests for materials and code should be addressed to corresponding authors.

## Supplementary Materials

### Materials and Methods

#### Tadpole generation and husbandry

Tadpoles were generated and staged as previously described (*13*). After NF Stage 45, tadpoles were fed once or twice a day with filamentous blue-green algae (ZM spirulina powder) suspended in water. Wild-type *Xenopus laevis* were used for experiments unless otherwise stated. Tadpoles classified as regeneration-competent were NF Stage 52-53, regeneration-restricted were NF 55-56, and regeneration-incompetent were NF Stage 58-60. Animal experiments were approved by the University Biomedical Services at the University of Cambridge and complied with UK Home Office guidelines (Animal Act 1986).

#### Single-cell dissociation, library preparation and sequencing

For developmental samples, tadpoles were killed, and samples were collected at the aforementioned stages. For amputation/regeneration samples, tadpoles were anaesthetized by incubating them with 0.1X MMR 0.002% MS222 (A0377876, Acros Organics), placed on a wet towel and the right hindlimbs were amputated at the presumptive knee/ankle level for regeneration-competent tadpoles, and at the ankle level for –restricted or –incompetent tadpoles. Afterwards, the tadpoles were returned to fresh water. At 5 days post amputation (dpa), tadpoles were killed and the newly generated tissues on the amputation plane were collected. Contralateral control samples were also collected from these tadpoles, and intact limb buds or autopods including ankle were collected. For each scRNA-Seq experiment, tissues were collected from a total of 8-10 tadpoles to reduce variance caused by staging differences. Dissociations were performed on a pool of 4 limbs in an Eppendorf tube with the following protocol. First the samples were washed with Ca-Mg free 1X MBS ((Barth-HEPES Saline) 10X stock: 88 mM NaCl, 1 mM KCl, 2,4 mM NaHCO3, 0.82mM MgSO4.7H2O, 0.33mM Ca(NO3)2.4H2O, 0.41 mM Cacl2.6H2O, 10 mM HEPES. Add ∼3 mL of 10N NaOH to obtain a pH of 7.4 to 7.6). Samples were then incubated with 1X Trypsin (Sigma, 59427C) in Ca-Mg free 1X MBS with 0.5 µM EDTA for 10 minutes at room-temperature (RT) on a bench-top shaker at a speed of 300 rpm. Trypsin reaction was diluted with Ca-Mg free 1X MBS after 10 minutes. Physical dispersion was applied (10-15 times up-down trituration with a pipette) to samples before, half way, and at the end of trypsinisation. Cells were spun down at 250 g for 5 minutes, the supernatant was taken out, and cells were then resuspended in 1X Ca-Mg free 1X MBS. Cells were passed through a 35 µm diameter cell strainer then stained with 20 µM Hoechst 33342 (Sigma, 2261) in 1X Ca-Mg free MBS for 10-15 minutes, and Hoechst positive cells were sorted using a Sony SH800s Cell Sorter. scRNA-seq libraries were generated using 10X Genomics (v3 chemistry) and sequenced on an Illumina Novaseq 6000 SP flow cell.

#### scRNA-seq: data processing

Output files from 10X Genomics were processed using CellRanger v3.0.2, with sequences mapped to the *Xenopus* laevis 9.1 genome (Xenbase, ftp://ftp.xenbase.org/pub/Genomics/JGI/Xenla9.1/Xla.v91.repeatMasked.fa.gz and ftp://ftp.xenbase.org/pub/Genomics/JGI/Xenla9.1/1.8.3.2/XL_9.1_v1.8.3.2.allTranscripts.gff3.gz). Raw counts were normalized by cell library size, and then converted to TPX (transcripts per 10^4^). Cell calling was performed using CellRanger with default parameters. We further filtered the data according to library size, discarding cells with a total UMI count in the lowest quartile. We note that the main cell types and transcriptional changes remained unchanged if we omitted this cell-filtering step, although the clustering and visualization appears less robust (Fig. S4).

#### scRNA-seq: feature selection

Highly variable genes (HVGs) were selected for clustering and visualization as described previously (*13*) (Fano factor > 65^th^ percentile, mean expression > 5^th^ percentile and mean expression < 80^th^ percentile). Our initial analysis revealed that visualization and clustering was strongly influenced by cell cycle state (Fig. S2). To further refine the set of HVGs, we performed factor analysis with the aim of removing genes significantly associated with the cell cycle. Specifically, non-negative matrix factorization was performed on the cosine normalized, log2-transformed normalized counts matrix, using k = 30 components (R package *nnlm*). Factors were manually annotated according to their expression on the UMAP projection, and by inspection of the highest gene loadings for each factor; 2 factors corresponded to the cell cycle. To minimize the effect of the cell cycle signature on projection/clustering, we identified genes associated with these cell cycle factors (top 10% gene loadings for each factor) and removed these from the set of HVGs.

#### scRNA-seq: visualization and clustering

Data were projected onto two dimensions using the UMAP algorithm (*27*), with log2-transformed HVGs, cosine distance as a similarity measure, and parameters k = 15, min_dist = 0.2. Clustering was performed as described previously (*13*). Briefly, we constructed a graph using the UMAP function *fuzzy_simplicial_set* with k = 10 nearest-neighbors, and then performed graphical clustering using the walktrap algorithm (*cluster_walktrap* from R package *igraph*, with steps = 10).

#### scRNA-seq: gene set enrichment and cell cycle analysis

Single cell gene set enrichment scores were calculated with the *AUCell* R package (*28*), using HVGs as the background gene set. Cell cycle phase was inferred using *CellCycleScoring* (R package *Seurat*) (*29*).

#### scRNA-seq: annotation of cell-types

Cell type annotation was performed by manually comparing cluster-specific gene expression patterns (computed using *findMarkers* in R package *scran* (*30*)) with known cell type markers from the literature. Many clusters could be assigned to a well-characterized, functional cell type (e.g. *Satellite cell*). Other clusters could not be unambiguously identified, but were assigned a broad label together with a numeric identifier (e.g. *Blood 1*). Finally, a few clusters remain unannotated (e.g. *Unknown 1*). Dotplots of key marker genes of each cell type are provided in Fig. S5.

#### scRNA-seq: gene expression visualization

Gene expression in individual cells is visualized on the UMAP projection with points colored according to expression level (log10-transformed). Gene expression across groups of cells (e.g. for different clusters, or for different stage tadpoles) is shown using dotplots colored by mean expression (log10-transformed, normalized to group with maximal expression). We can detect alleles from both the Large (*Gene.L*) or Short (*Gene.S*) chromosomes present in the pseudotetraploid *Xenopus laevis* genome. In some figures, we report expression from both the large and short allele; in others, we report whichever allele has higher expression for brevity.

#### Regeneration assay and bead experiments

Affi-gel blue gel beads (Bio-rad, **1537301**) were incubated with 0.1% BSA or 1 µg recombinant human FGF10 (R&D, 345-FG) in 1-2 µl 0.1% BSA overnight at 4 degrees. Tadpoles were anaesthetized with 0.002% MS222, placed on a wet towel, and both right and left hindlimbs were amputated from ankle level in either –restricted or –incompetent tadpoles. 3-4 beads were placed on the amputation plane of the right hindlimb. Left hindlimbs served as an internal control for the experiments. Please note that pushing the bead deep in the tissues at the amputation site was avoided as much as possible, and beads were gently positioned instead. Tadpoles were monitored on a wet towel for 3-5 minutes then tadpoles that kept the beads were placed in fresh water. Tadpoles were killed in between 18-21 dpa to assess the regeneration outcome. The difference in the number of digits or digit-like structures between the right to the left limb was quantified for each tadpole.

#### Whole-mount mRNA visualisation, hybridization chain reaction (HCR), with or without combination of immunofluorescence or histology

##### HCR on whole limb or tail samples

HCR was applied as described before (*31*) with modifications, and materials for HCR were purchased from Molecular Instruments Inc unless otherwise stated. Limb and tail samples were fixed with 4% formaldehyde in 1X PBS for 40-60 minutes, permeabilized in 70% ethanol in 1X PBS for 2-4 hours, washed briefly with 1X PBS and collected in Eppendorf tubes. These procedures were carried out on a rotator at RT. The supernatant was taken out, 500 µl wash solution (Molecular Instruments Inc.) was added, and samples were rotated at RT for 5 minutes. The supernatant was taken out and replaced by 400-500 µl hybridization buffer (Molecular Instruments Inc.) for a 30 minutes incubation at 37 degrees. In parallel, the probe solution was prepared by diluting mRNAs targeting probes to 30-40 nM in 200 µl hybridization buffer and incubated for 30 min at 37 degrees. The hybridization buffer from samples were taken out and probe solution was placed on samples for a 12-16 hours incubation at 37 degrees. Subsequently, the samples were washed 2 × 20 minutes with wash buffer, and 2×30 minutes with 5x SSC-T at RT. To visualize probes, amplification solution was prepared by first heating to 95 degrees for 90 seconds the fluorophore attached hairpins pairs (h1 and h2 hairpins) that matches to the probes. Hairpins were then left in dark at RT for 30 minutes. Afterwards, final amplification solution was prepared at 40-60 nM h1 and h2 in 200 µl amplification buffer. Afterwards, samples were placed in amplification solution at room temperature, protected from light, for 12-16 hours on a rotator. Samples were washed with 2×20 min SSC-T. Samples were then put in 1X PBS.

##### Whole-mount HCR samples imaging

For stereomicroscope or confocal imaging of whole samples, the samples were mounted in 0.6%-0.8% ultra-low gelling temperature agar (Sigma, A5030) in 1X PBS.

##### Sectioning of samples after HCR

In the subsequent step of the protocol, the samples were protected from light to preserve the HCR signal. The samples were incubated in 15% sucrose in 1X PBS at RT for 1 hour, then 30% sucrose in 1X PBS at 4 degrees overnight. Samples were then placed in O.C.T. solution and incubated at −80 overnight. Samples were cryosectioned to 5 µm thickness, stained with 20 µM Hoechst (Sigma, 2261) in 1X PBS at RT for 10 minutes and imaged.

##### Immunostaining

After sectioning of HCR stained limb, the samples were processed for immunostaining. Samples were blocked with 50% Cas-Block (Invitrogen, 008120) in 1X PBS-T (1X PBS + 0.1 Tween-100) and incubated for 30 minutes in room temperature without rotating. Samples were then incubated with antibodies (listed below) at 4 degrees overnight without rotating. Samples were washed with PBS-T for 2×10 minutes, blocked by 50% Cas-Block in 1X PBS-T for 30 minutes, and incubated with secondary antibodies (listed below) for 1 hour, all these steps were carried out at RT without rotating. Samples were washed with 1X PBS-T for 2×10 minutes and 2×20 minutes 1X PBS at RT without rotating. After antibody staining, samples were stained with Hoechst and washed with 1×5 min 1X PBS at RT without rotating. Samples were mounted in 80% Glycerol in 1X PBS with a coverslip and imaged.

Tail whole-mount HCR staining can be combined with whole-mount immunofluorescence by following the above immunofluorescence protocol except that the mounting of whole-tails were done in ultra-low gelling temperature agar for imaging.

HCR probes and Hairpins: Probes for *Fgf8.L, Dpt.L, Htra3.L, Prrx1.L* and *Sp9.L* were purchased from Molecular Instruments Inc.. Probes were designed against the full-length *Xenopus Lgr5.S, Msx1.L*, and *Fgf10.L* mRNA sequence as described by (*32*). HCR Hairpins were purchased from Molecular Instruments Inc.

Primary antibodies, and working dilutions: TP63 [4A4] (Abcam, ab735, 1:200), B-CATENIN (Abcam, ab6302, 1:2000), E-CADHERIN (5D3, DSHB, 1:10), ITGB1 (8C8, DSHB, 1:10), anti-EGFP (Abcam, ab13970, 1:500).

Secondary antibodies: goat anti-chicken IgY (H+L) secondary antibody, Alexa Fluor 488 (Invitrogen, A11039, 1:500), goat anti-mouse IgG (H+L) cross-adsorbed ReadyProbes secondary antibody, Alexa Fluor 594 (Invitrogen, R37121, 1:500). goat anti-mouse IgG (H+L) cross-adsorbed ReadyProbes secondary antibody, Alexa Fluor 488 (Invitrogen, R37120, 1:500).

Leica SP8 upright confocal microscope with a 40x/1.3 HC PL Apo CS2 Oil objective was used for all confocal images except for Fig. S8B images which were taken with Leica SP8 inverted confocal microscope with a 20x/0.75 HC PL Apo CS2 Multi. LAS X was used for setting tiled images, and 20% overlap between tiles were used. Limb whole-mount HCR images were taken via a Leica stereomicroscope equipped with a DFC7000 T camera. Fiji was used for maximum projection of z-stacks and to adjust contrast to highlight biological relevance. If needed, images were cropped, flipped, and/or rotated to highlight biological relevance.

Histological staining can be done on top of cryosectioned HCR samples. Briefly, samples were stained with hematoxylin and eosin according to manufacturer’s protocol (Abcam, ab245880), afterwards samples were stained for Alcian Blue (Sigma, B8438) according to manufacturer’s protocol. Histology images were taken on a Zeiss AxioImager compound microscope.

#### Ex vivo limb culture method to assess AER cell formation and proximal chondrogenesis

Limbs were first amputated from presumptive knee/ankle level for – competent and ankle level for –restricted or –incompetent tadpoles. The distal parts of these amputated explants were then removed and the remaining proximal segment was placed in 1000, 500, 200 µl explant media (L-15 (Thermo, 11415064) 1X Antibiotic-Antimycotic (Thermo, 15240062), 20% Fetal Bovine Serum Superior (Sigma, S0615)) in 12, 24, or 96-well plates, respectively. Explants were cultured for 3 days without changing the media. After 3 days, to quantify AER cell formation the explants were fixed and proceeded to HCR protocol; to quantify proximal chondrogenesis the explants were fixed with 4% formaldehyde, mounted in 0.6% Low-Melt agar, and directly imaged via Stereomicroscopy. Explants emit autofluorescence. Though the abundant HCR signal can be seen despite the autofluorescence, to discriminate the HCR signal from autofluorescence in finer detail, sample images were taken in red and green channel separately with the same exposure and gain settings, and then merged in Fiji. In merged images, the background signal due to autofluorescence was visualized as yellow and the HCR signal was either red or green. As AER cells were largely detected as a monolayer population, AER cell formation was calculated by measuring the length of the *Fgf8.L* signal on the amputation plane using Fiji segmented line option. The proximal chondrogenesis can be visually distinguished, and to determine the chondrogenesis length, chondrogenic structure length from top to bottom was also measured using Fiji. Samples where a clear chondrogenesis was not visible were omitted from further analysis. These images were taken in brightfield imaging and measurements were done in Fiji.

For drug and recombinant protein treatments, the explants were placed in culture media containing the following small molecules concentration or recombinant protein amounts, unless otherwise stated: 100 µM ICRT3 (Sigma, SML0211), 100 µM SU-5402 (Sigma, SML0443), 50 µM SB-505124 (Sigma, S4696), 100 µM DAPT (Sigma, D5942), 2.5 µM LDN-193189 (Stemgent, 04-0074), 500 ng human recombinant FGF10 (R&D, 345-FG), 1.25 µg human recombinant NOGGIN (R&D 6057-NG), and 500 ng human recombinant BMP4 (R&D, 314-BP). Drugs were prepared in DMSO, and recombinant proteins were prepared in 0.1% BSA. Small molecule experiments were conducted in 24-well plate. Recombinant protein experiments were done in 96-well plate. Max 5-6 explants were placed in 24-well plates. 1 explant was put in one well of 96-well plate for recombinant protein treatments. In all chemical and recombinant protein perturbation experiments, one limb of the same animal was subjected to the perturbation, and the contralateral limb served as a control. These control explants were exposed to solution containing matching DMSO or BSA concentration in 1X PBS for chemical or recombinant protein perturbations, respectively. Perturbation and control samples were pooled separately at the end of experiments and proceeded with staining.

#### EdU Labelling

*Ex vivo* limbs were cultured with 10 µM EdU (Thermo, C10337) for 3 days in dark foiled cover. Afterwards, samples were fixed, and *Fgf8.L* mRNA was stained using the HCR protocol, followed by cryosectionning, as described above. Sections were subjected to Click-It reaction as described in manufacturer’s protocol (Thermo, C10337). Hoechst was added at the end of the protocol. Samples were visualized by confocal microscopy as described above. (1) *Fgf8.L* positive cells, and (2) EdU positive and *Fgf8.L* positive cells on the amputation plane were manually counted, and the percentage of EdU positive *Fgf8.L* positive cells were calculated for each sample.

#### Bead experiment for proximal chondrogenesis

Beads were prepared as described above. Explants from –restricted tadpoles were harvested as described above and beads were implanted on the proximal site of explants. At 3 dpa, explants that did not contain bead at their proximal site anymore (presumably due to repelling) were omitted from further analysis. At 3 dpa, samples were imaged without fixation and the extent of chondrogenesis was measured by Fiji.

#### DiO Labelling

DiO (DiO’; DiOC_18_(3) (3,3’Dioctadecyloxacarbocyanine Perchlorate), Thermo, D275) was prepared by dipping a tip in the DiO containing powder tube, and placing the tip in a 10 µl 100% ethanol containing Eppendorf. A glass needle tip was then dipped in the diluted DiO solution and harvested *ex vivo* limbs were labelled on a wet towel. These cultures were placed in *ex vivo* culture media and explants were imaged every day with a stereomicroscope.

#### Ex vivo limb co-culture, and conditioned media experiments

For co-culture experiments, one -competent and one –incompetent limb explants were incubated together in 200 µl explant media in a well of 96-well plate. For antibody experiments, one limb of each animal served as a control and was incubated with 1 µg Rabbit-IGG isotype control antibody (ab37415) while the contralateral limb was incubated with 1 µg anti-NOGGIN antibody (ab16054). Antibodies and media were only added at the beginning of the cultures and were not replaced during the experiment.

For conditioned media experiments, conditioned media supplying and receiving explants were prepared separately. Supplying explants were prepared one day before harvesting receiving explants and incubated in 200 µl explant media in a well of 96-well plate. After one day, media from the supplying explant was collected and used to culture the newly harvested receiving explant, and a fresh media was added for supplying explant. This change of media procedure was repeated for 3 days. For antibody experiments, supplying explant media was collected and pre-incubated with 1 µg antibodies for 25-30 minutes at RT on a rotator, then the pre-incubated media was placed on the receiving explants.

#### Replicate information and statistical tests

Sample sizes were not pre-determined in any experimental setup. In this work, biological replicates refer to samples obtained from multiple animal batches and to experiments carried out different days. In all experiments, wild-type tadpoles were used from tanks that contain multiple batches (tadpoles raised from different father and/or mother). In all explant perturbation experiments, samples were compared to their contralateral controls, and a Mann Whitney U test was used to determine statistical significance. For regeneration and bead experiments, t-test was used.

#### Data availability

Code is available at https://github.com/MarioniLab/XenopusLimbRegeneration2020. Sequencing data, together with processed counts matrices, are available on ArrayExpress with the accession number E-MTAB-9104. We provide an interactive online tool to explore our dataset https://marionilab.cruk.cam.ac.uk/XenopusLimbRegeneration/

---

**Fig S1:**
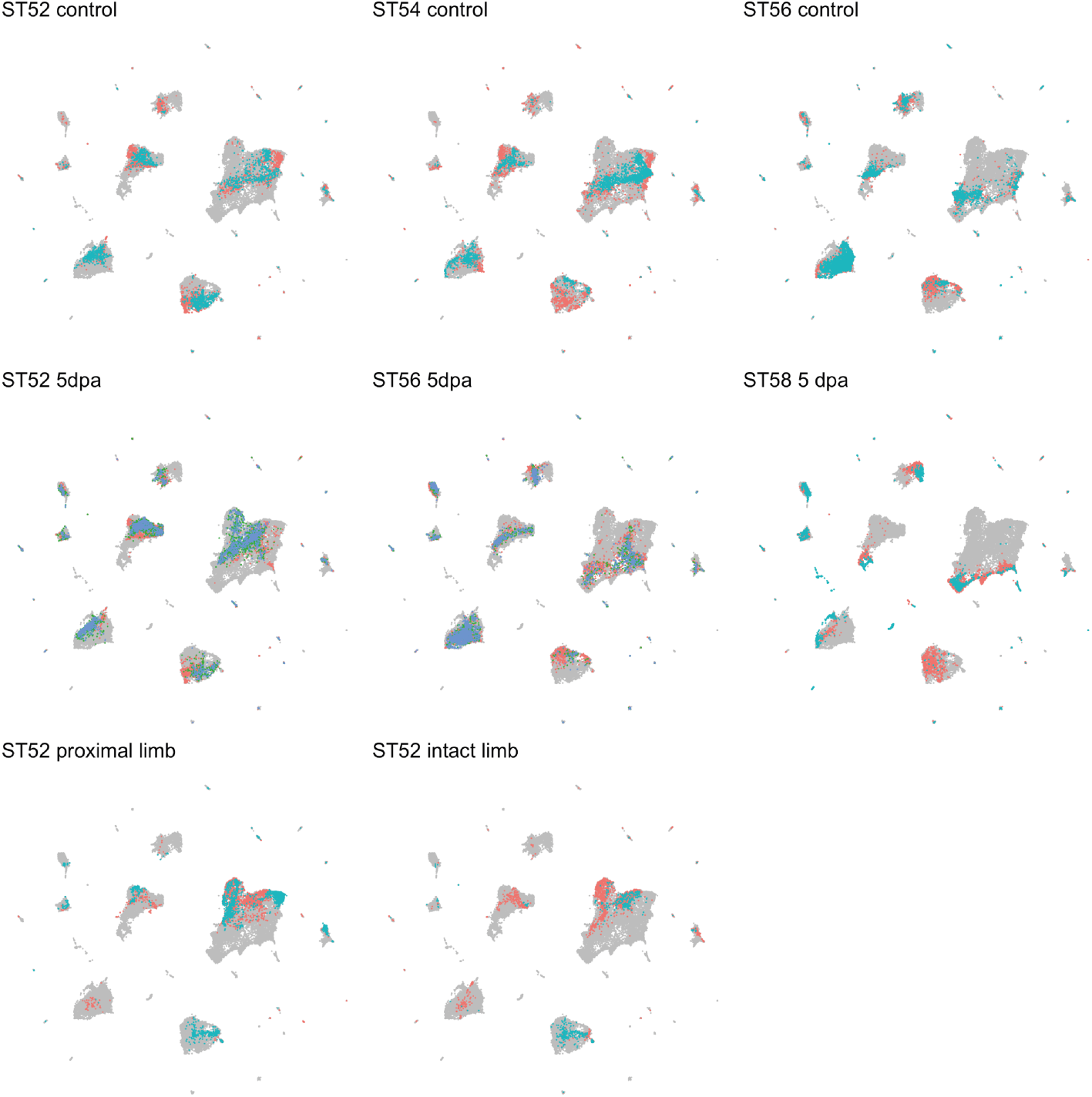
Contribution of different conditions to the pooled UMAP projection. UMAP visualization of cells from all conditions and replicates, allowing the identification of transcriptional changes that are consistent across replicates. Grey dots: cells from all samples; red, blue, green dots: cells from different biological replicates for the selected sample.

**Fig S2:**
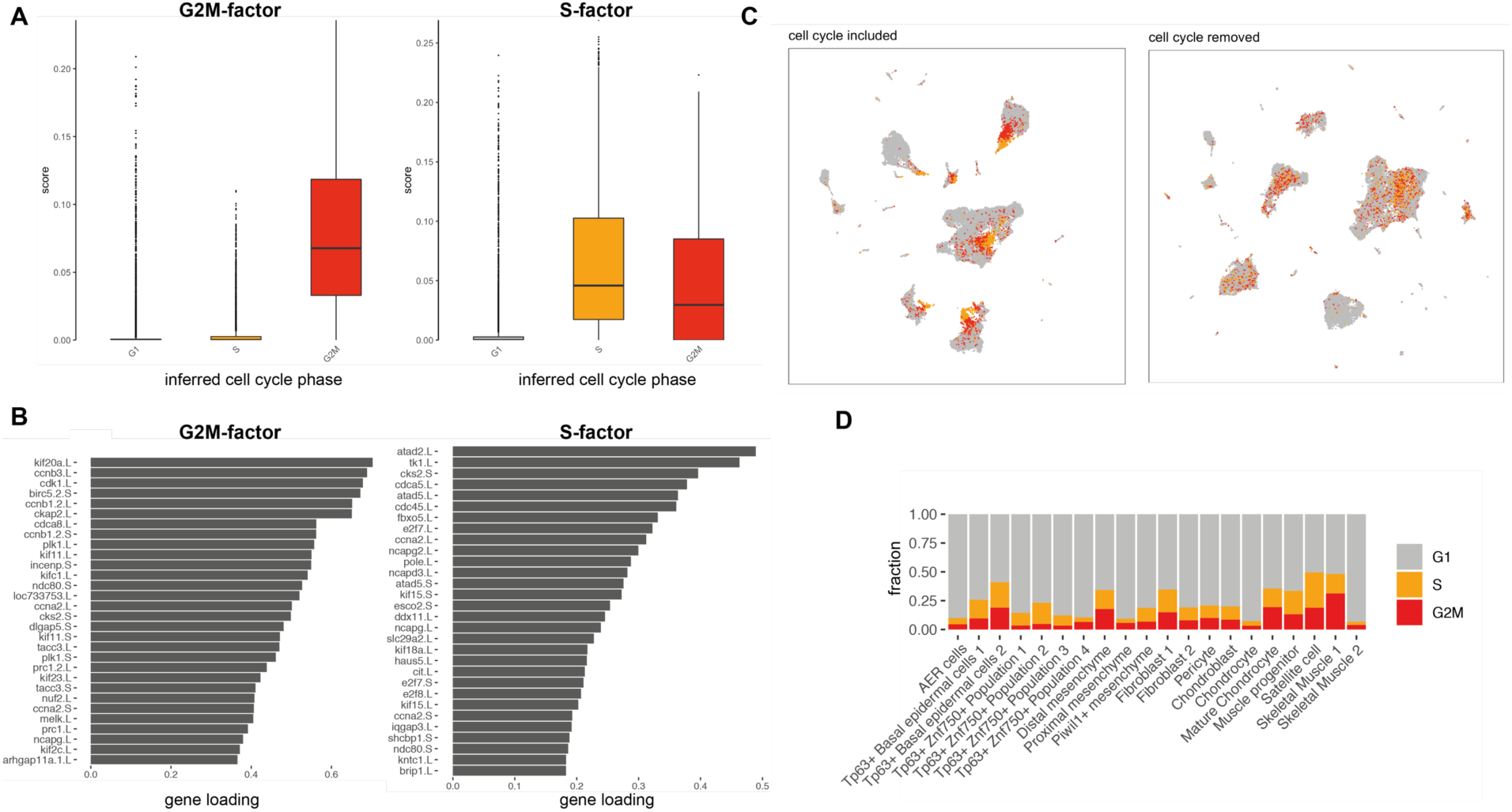
Detection and removal of the cell cycle signature. **A)** Unbiased factor analysis identified two factors that correspond to computationally-inferred cell cycle phases (G2M-factor, left; S-factor, right). **B)** Factor loadings for the top 30 genes associated with cell cycle factors. **C)** Removal of genes with high loadings for either G2M- or S-factors significantly reduces the influence of cell cycle phase on the UMAP projection. Dot colour indicates inferred cell cycle phase. **D)** Inferred cell cycle states for selected cell types.

**Fig S3.**
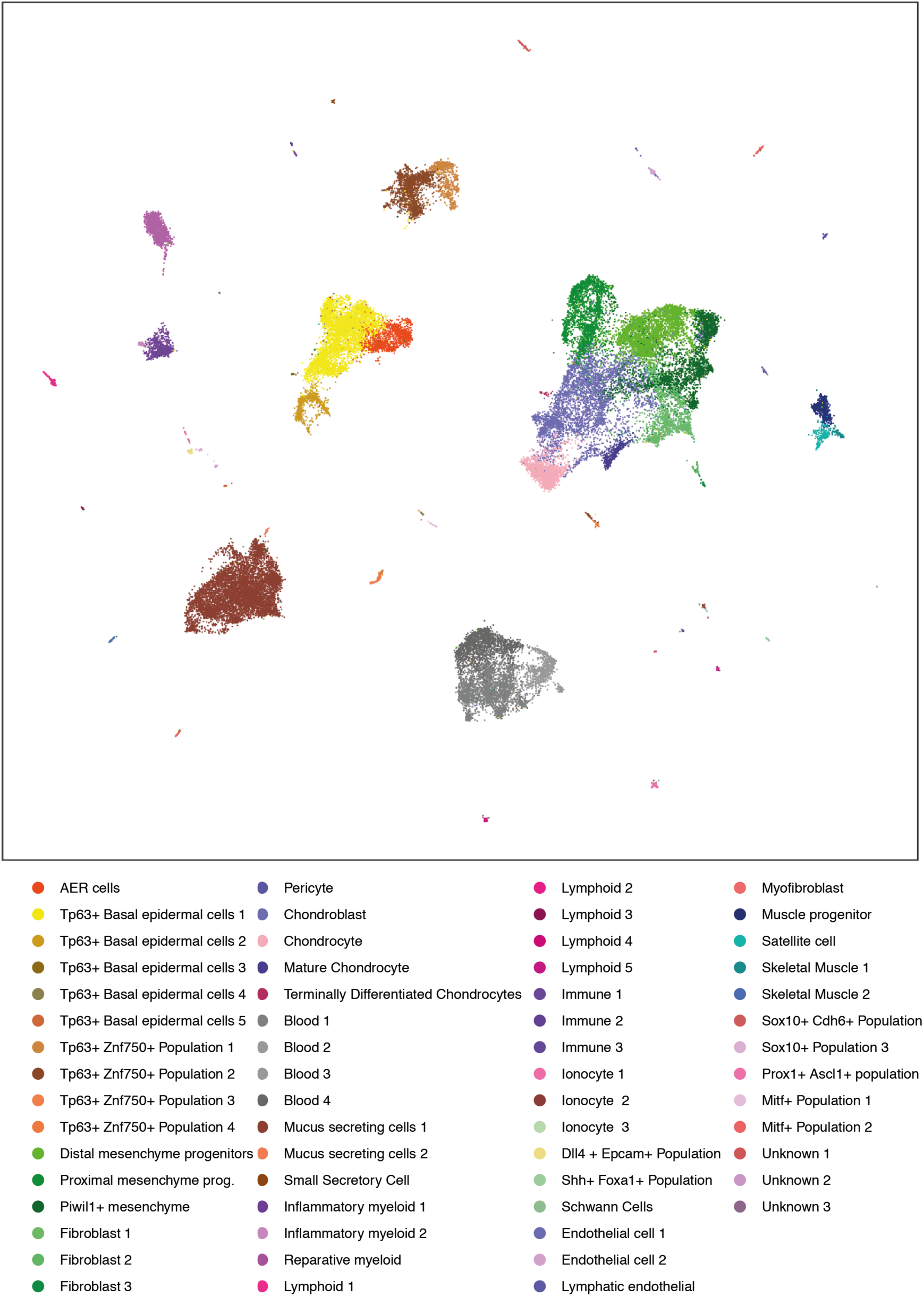
An atlas of cell types in developing and amputated limbs at different stages of regeneration-competence. Pooled UMAP visualization of *Xenopus* limb cells, with colours representing distinct cluster identities.

**Fig S4.**
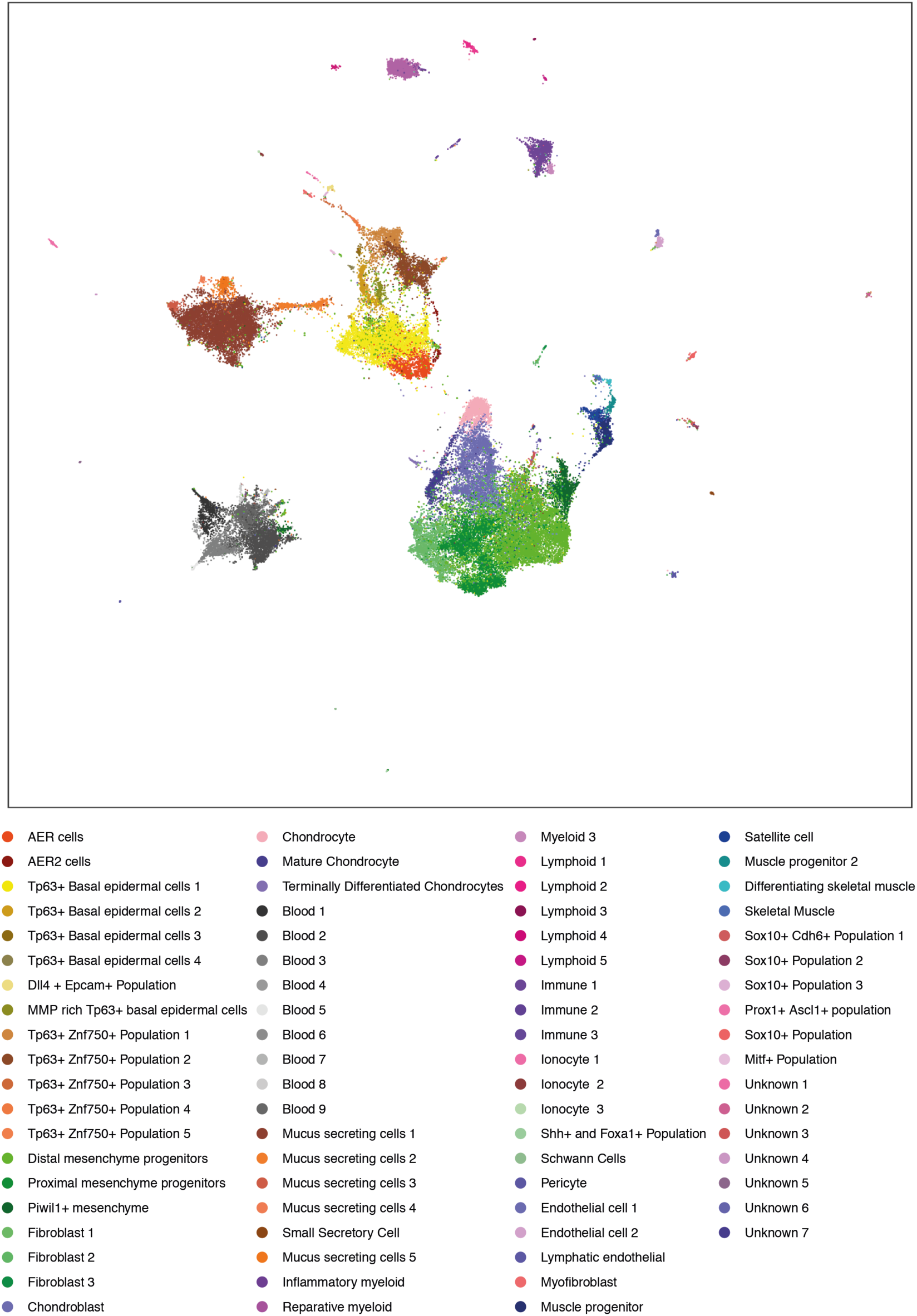
An expanded atlas of the *Xenopus* limb using less stringent cell filtering protocols. Pooled UMAP visualization and clustering of all barcodes that are identified as cells using cellRanger with default parameters. The majority of transcriptional states are similar to Fig S3, although a fraction of low-UMI mesenchymal cells appear mislocalized across the atlas.

**Fig S5:**
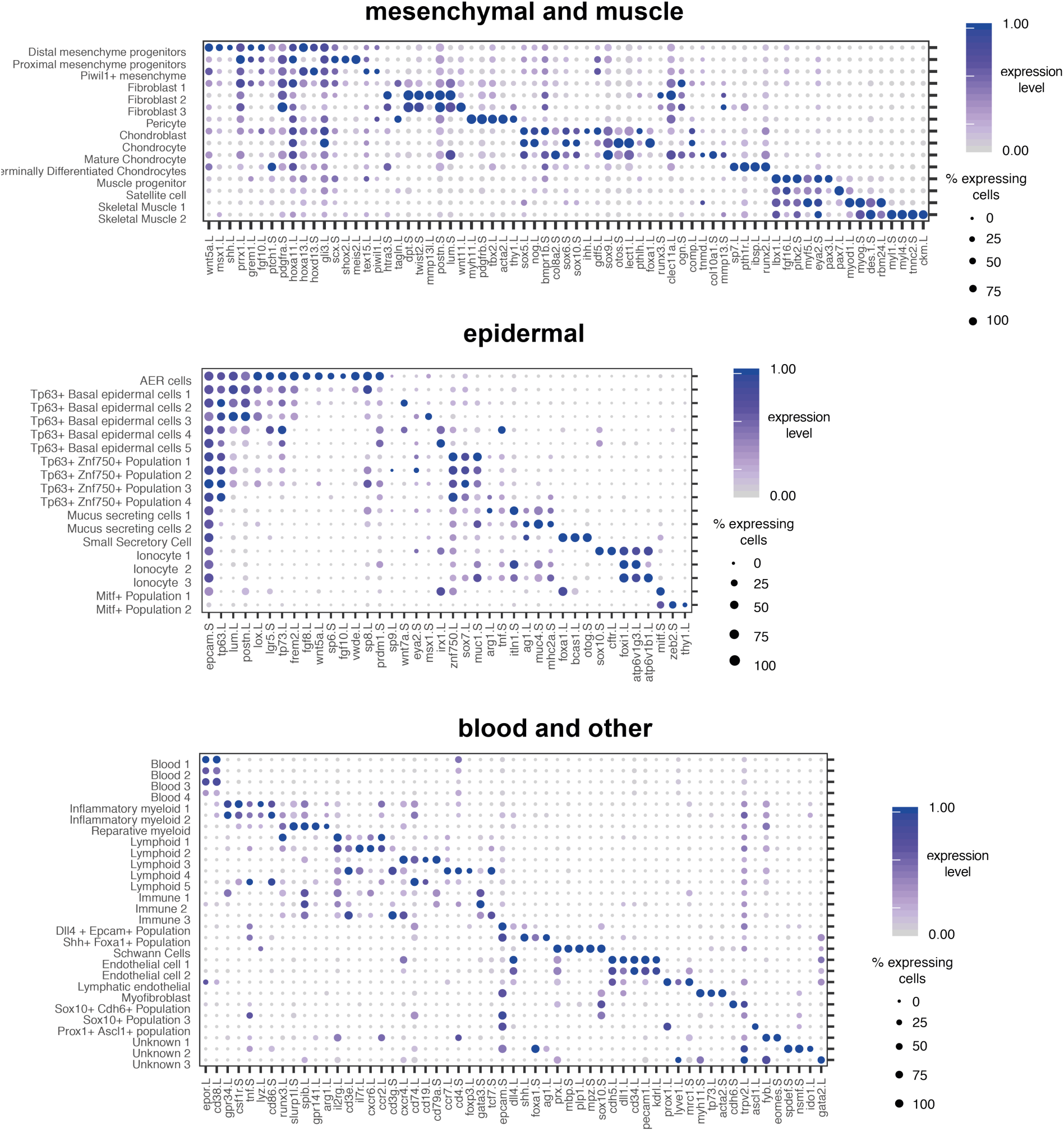
Annotation of cell types using known markers of cell identity. Dotplots showing marker genes for each of the 61 cell types in our atlas. For ease of presentation, we group cell types into three broad categories: mesenchymal and muscle (top), epidermal (middle), blood and other (bottom). Dot colour denotes mean expression level within the cluster; dot size denotes the percentage of cells within the cluster with non-zero expression.

**Fig S6.**
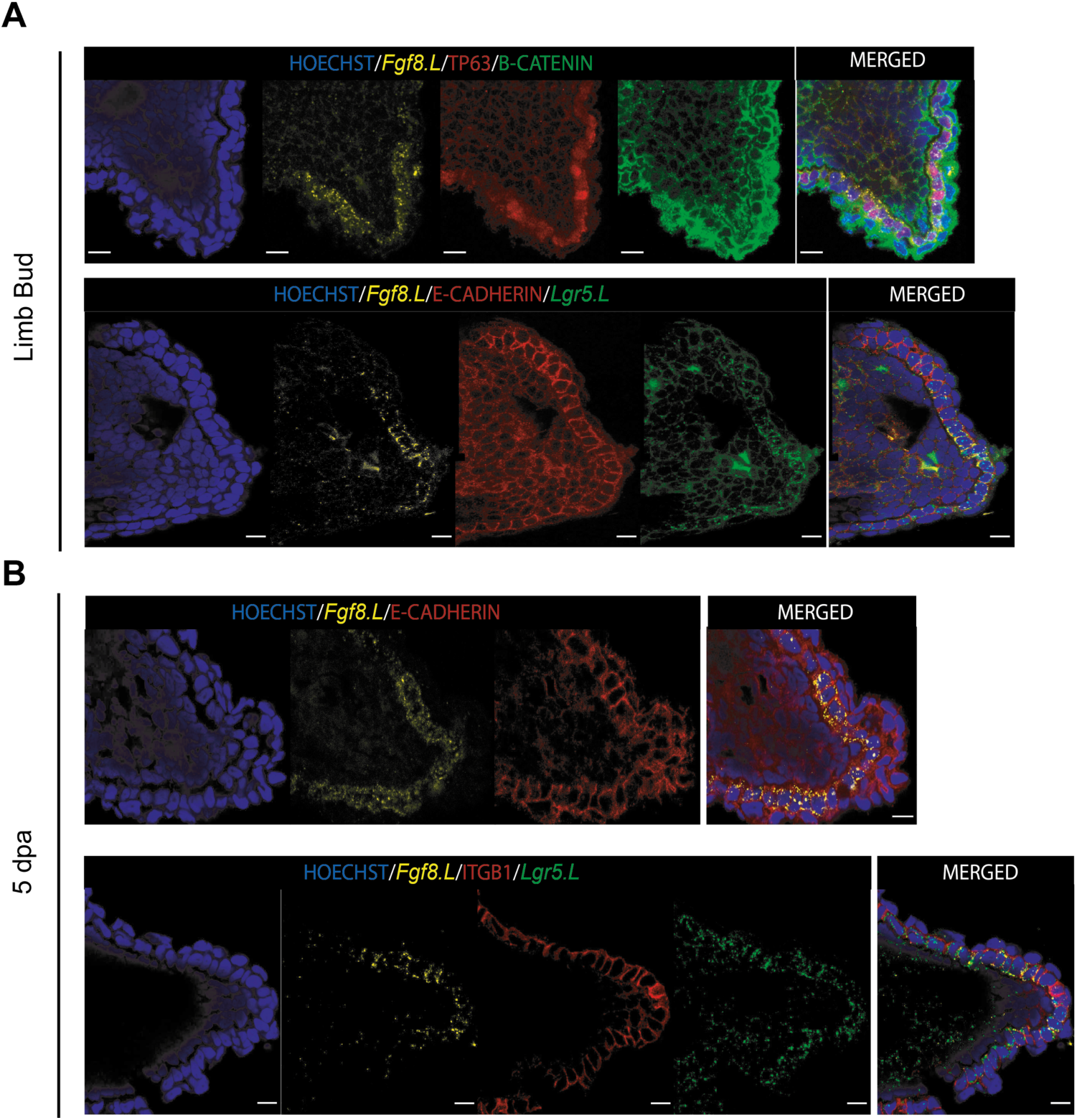
AER cells are largely found as cuboidal monolayer cells showing apical-basal polarity. AER cells were visualised during limb development **(A)** and at 5 dpa **(B)** in regeneration-competent tadpoles by labelling *Fgf8.L* mRNA. AER cells are largely present as monolayer cuboidal basal epidermal cells with apical-basal polarity. A simple squamous layer is present above AER cells, and cells with mesenchymal morphology are located underneath AER cells. From the proximal to distal midline of the epidermis, *Lgr5.S* expression is first detected, followed by *Fgf8.L* mRNA expression. Both *Fgf8.L* and *Lgr5.S* are expressed at high levels at the tip of limbs. AER cells show similar cuboidal morphology during development and regeneration. Basal epidermal cells are morphologically similar based on Hoechst and membrane markers, and *Fgf8.L* detection is required to detect AER cell. Row 1: Blue, Hoechst; Yellow, *Fgf8.L* mRNA; Red, TP63; Green, B-catenin. Row 2: Blue, Hoechst; Yellow, *Fgf8.L* mRNA; Red, E-Cadherin; Green, *Lgr5.S* mRNA. Row 3: Blue, Hoechst; Yellow, *Fgf8.L* mRNA; Red, E-Cadherin. Row 4: Blue, Hoechst; Yellow, *Fgf8.L* mRNA; Red, ITGB1; Green, *Lgr5.S* mRNA. Scale bars = 10 µm.

**Fig S7.**
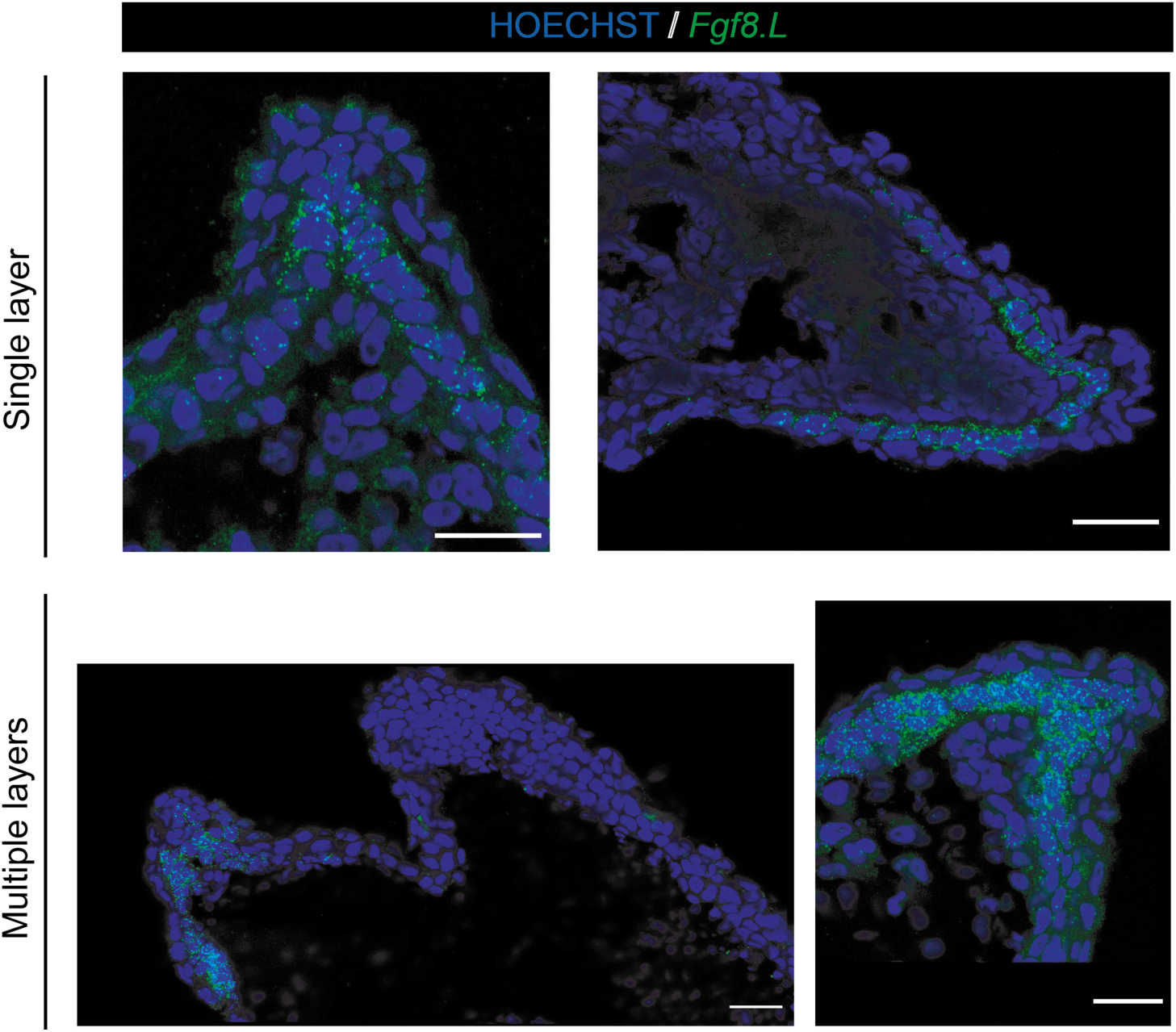
AER cells can be mono- or multi-layered structures. *Fgf8.L* images of sectioned 5 dpa samples from regeneration –competent (top) and –restricted (bottom) samples. Morphology of AER cells (*Fgf8.L+*) can vary between sections and samples. Top left, AER cells are seen as single monolayer largely cuboidal although some have higher height to width ratio. Top right, AER cells are seen as single monolayer largely cuboidal cells. Bottom left, AER cells can be seen as multi-layered population that is not covering the whole amputation plane. Bottom right, AER cells can be seen as multi-layered population covering the amputation plane. Blue, Hoechst; Green, *Fgf8.L* mRNA. Scale bars = 25 µm.

**Fig S8.**
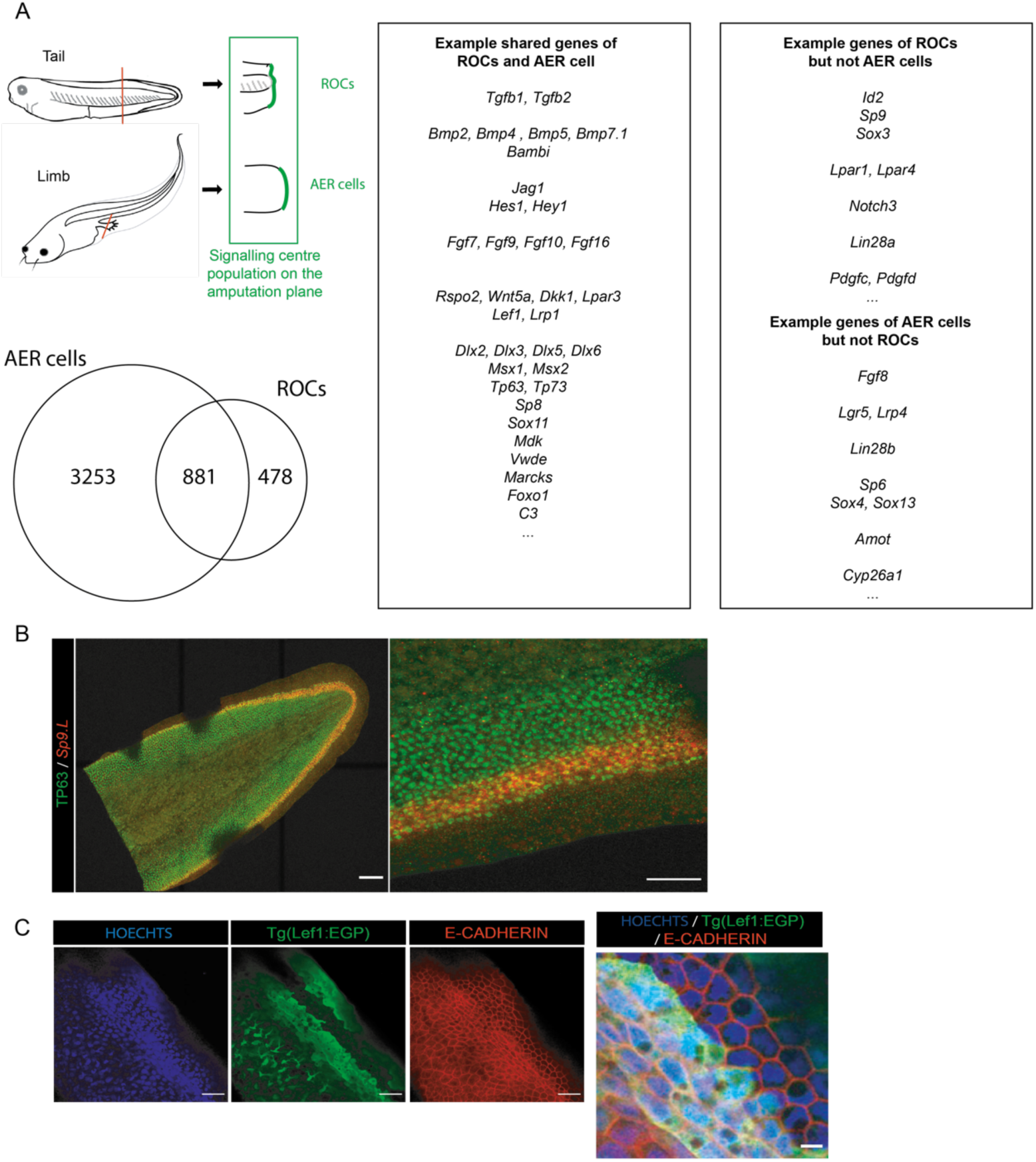
Specialised wound epidermis of tail and limb regeneration share some transcriptional similarities while presenting different cellular morphology. (Left) A signalling centre population serving as the specialised wound epidermis is associated with *Xenopus* tail and limb regeneration. However, tail uses regeneration-organising-cells (ROCs) (*13*) while limb uses AER cells for this purpose. Both AER cells and ROCs share the expression of many genes highlighting their similarity, although there are some genes that are unique to each population. AER-and ROC-specific genes were identified as genes significantly upregulated relative to other basal epidermal cells. (Right) A select number of genes, specifically ligands and transcription factors that are associated with regeneration, are highlighted. **(B)** ROCs and AER cells show different morphologies (please see Fig. S6 for AER cells). ROCs were visualized by staining NF Stage 40 by *Sp9.L* mRNA expression (highly specific for ROCs (*13*)) and TP63 immunolabelling for whole tail (Left) and zoomed in version (Right). In the zoomed in version for staining *Sp9.L* shows two level of expression in ROCs: a single outer layer of *Sp9.L* low cells, and multiple inner layers of Sp9.L high cells. Please note that this is not whole bottom-top image of a tail as evidenced by absence of TP63 staining in the in middle part of the tissue. Red, *Sp9.L* mRNA; Green, TP63. Scale bars= (left) 250 µm, (right) 100 µm. **(C)** ROCs are visualized using the pbin7LEF:GFP line, as defined previously (*13*), and E-CADHERIN staining was used to delineate cell shape. Inner layers of ROCs have flattened cell shape while the outside layer ROCs exhibit more square-like shape. ROCs do not have branched nuclei, unlike fin cells. Blue, Hoechst; Green, EGFP; Red, E-cadherin. Scale bars= 10 µm.

**Fig S9.**
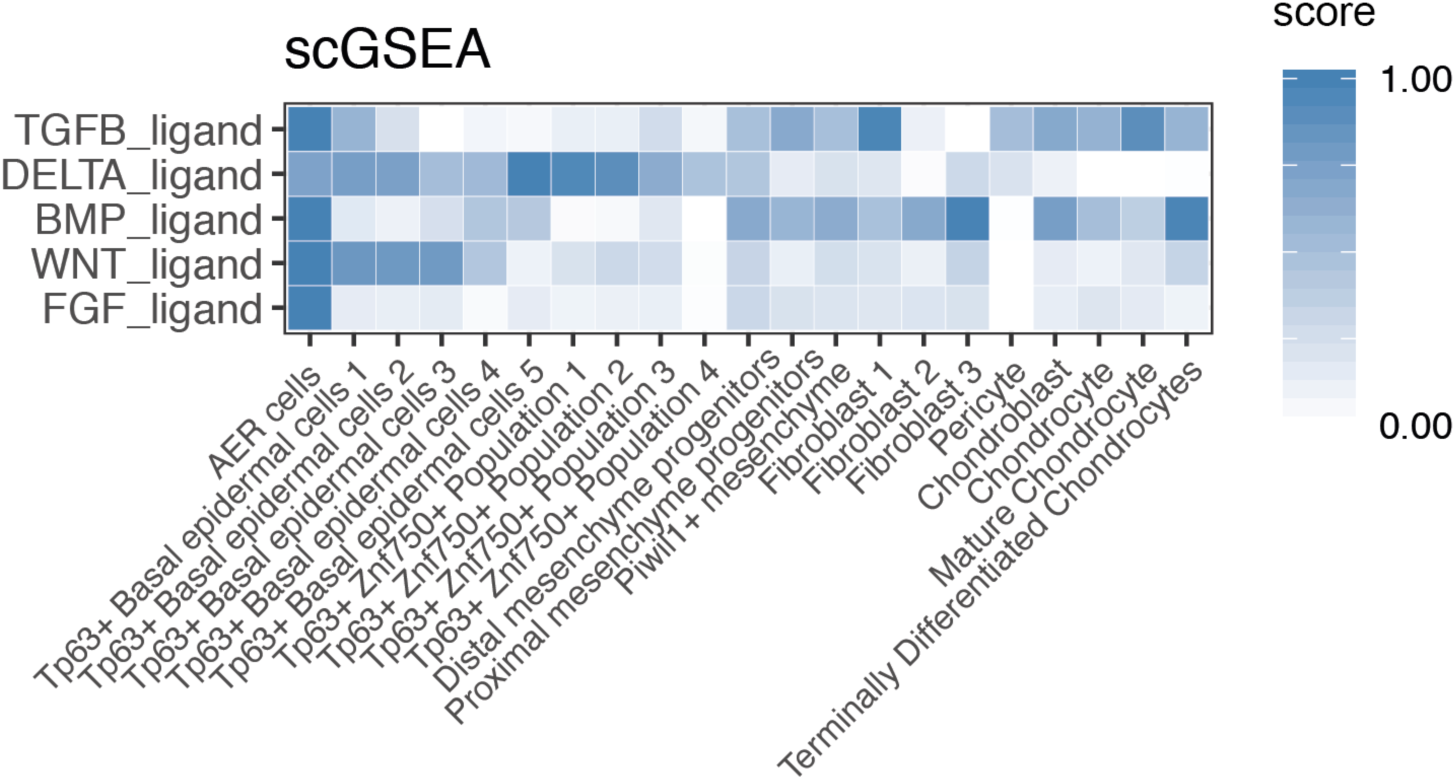
AER cells are a signalling centre population. Heatmap showing single-cell gene enrichment scores for ligands from the main signaling pathways are shown for epidermal cell types. AER cells have high signal center properties as they express high levels of TGF-β, Delta, BMP, WNT, and FGF ligands.

**Fig S10.**
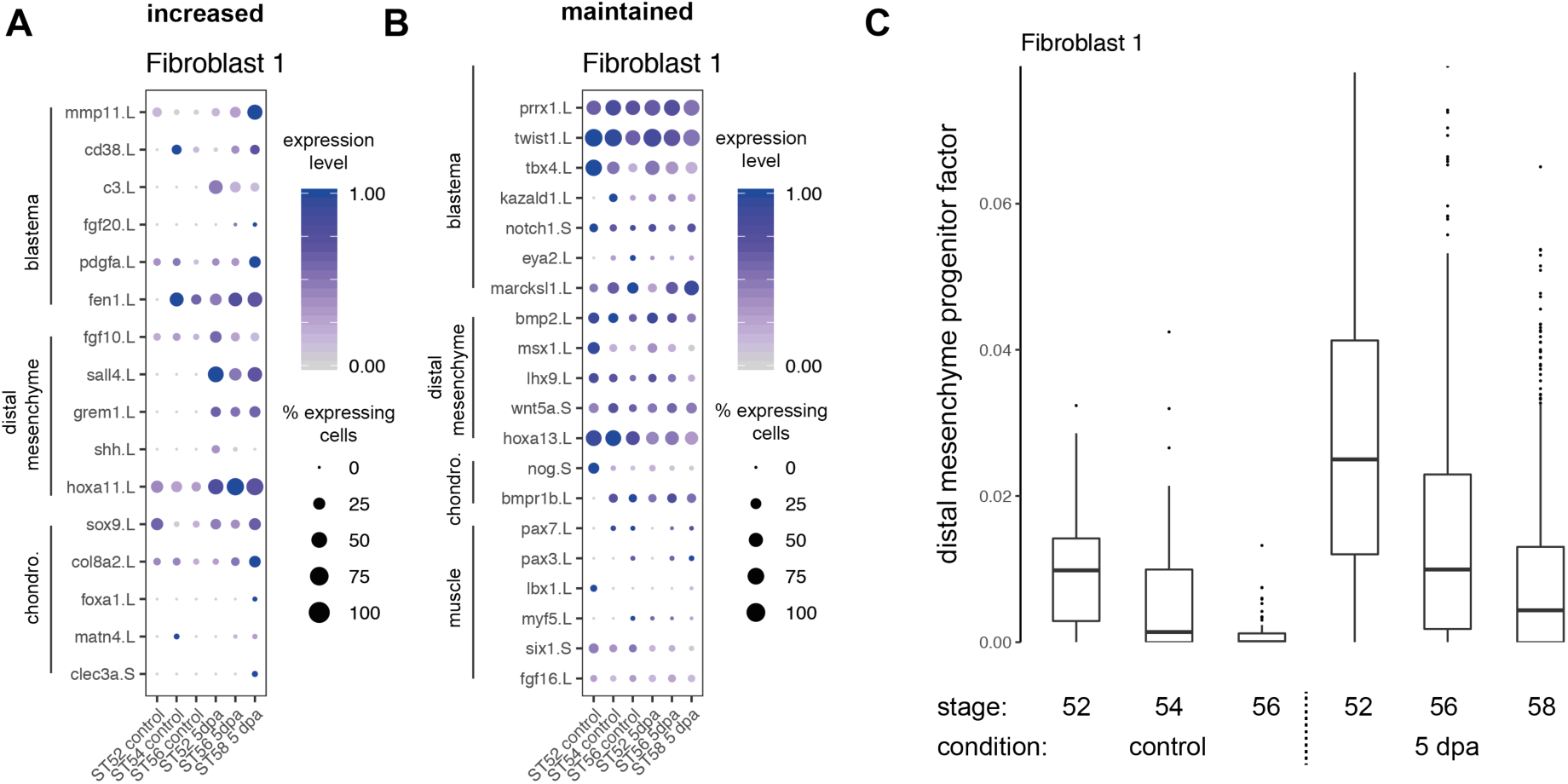
A subset of fibroblasts express dedifferentiation and blastema genes independently of the regeneration-outcome. Expression of genes and putative gene sets associated with regeneration in the Fibroblast 1 cluster, visualized using dotplots and factor analysis. **(A)** Expression of specific genes that increase upon amputation regardless of stage. **(B)** Expression of specific genes that are expressed in intact limbs and are maintained after injury. (**C)** Following amputation, the distal mesenchyme factor increases in Fibroblast 1 cells across all stages.

**Fig S11.**
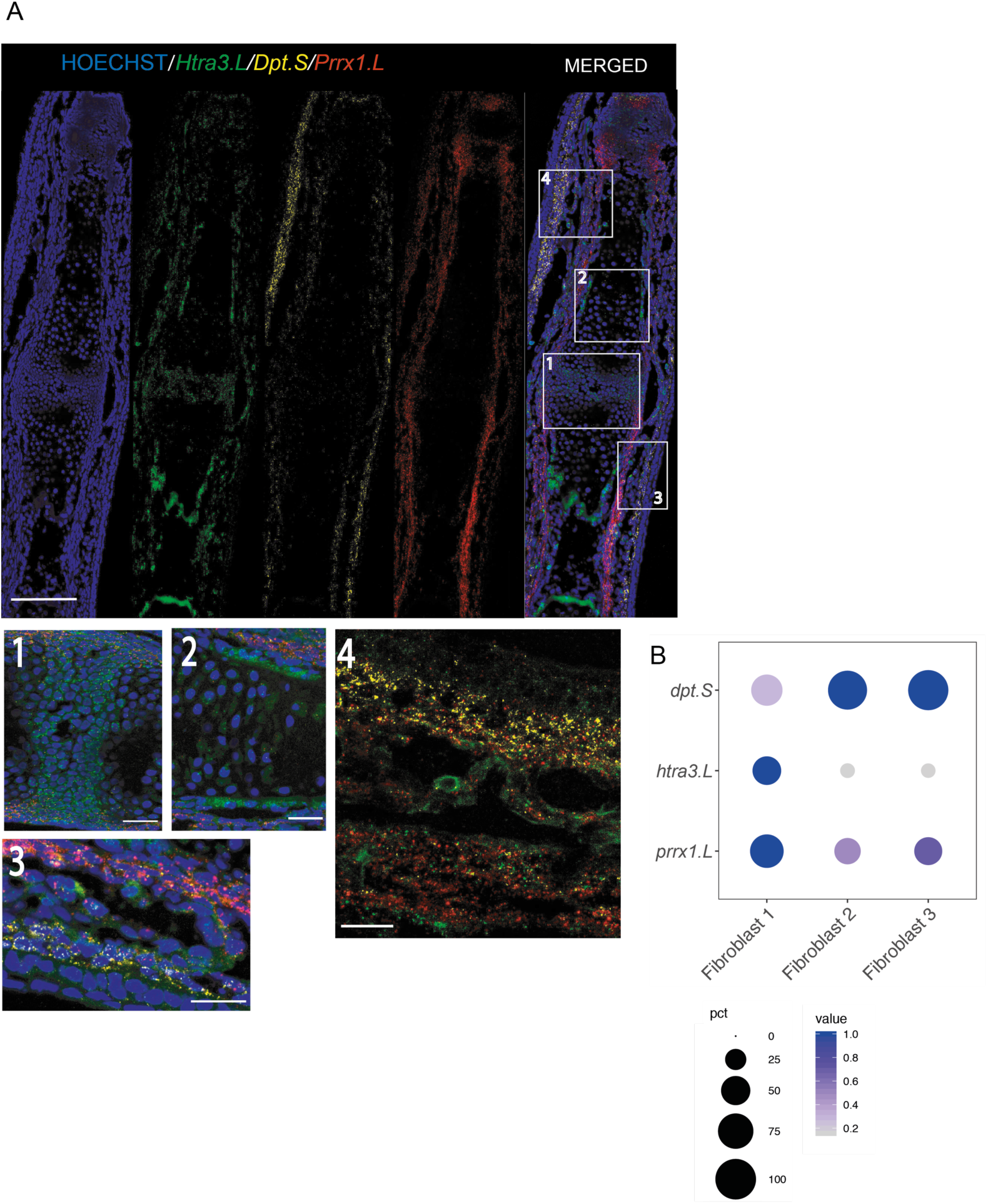
Fibroblast 1 cluster cells are largely found beneath skin cells and nearby perichondrial cells. **(A)** (Top) Confocal images of a Stage 56 digit stained against *Htra3.L, Prrx1.L*, and *Dpt.S.* Cells expressing *Htra3.L*/*Prrx1.L*/*Dpt.S* are found underneath the skin regions and nearby perichondrium regions. (Bottom) Zoomed in version of selected areas show: (1) joint forming regions are enriched for *Htra3.L* expression; (2) Inner perichondrial regions are enriched for *Htra3.L* and outer perichondrial regions are enriched for *Prrx1.L* expression. (3-4) Outerlayers of dermal fibroblast area enriched for *Dpt.S* and lower levels of *Prrx1.L* and *Htra3.L*. Inner layers of dermal fibroblasts/nearby perichondrial regions are enriched for higher Prrx1.L and lower *Dpt.S* and *Htra3.L* expressions. Blue, Hoechst; Green, *Htra3.L* mRNA; Red, *Prrx1.L* mRNA; Yellow, *Dpt.S* mRNA. Scale= 125 µm for top images, 25 µm for bottom no 1-3, and 20 µm for bottom no 4. **(B)** Dot plot showing expression of *Htra3.L, Prrx1.L*, and *Dpt.S* for Fibroblast 1, 2, and 3 clusters.

**Fig S12.**
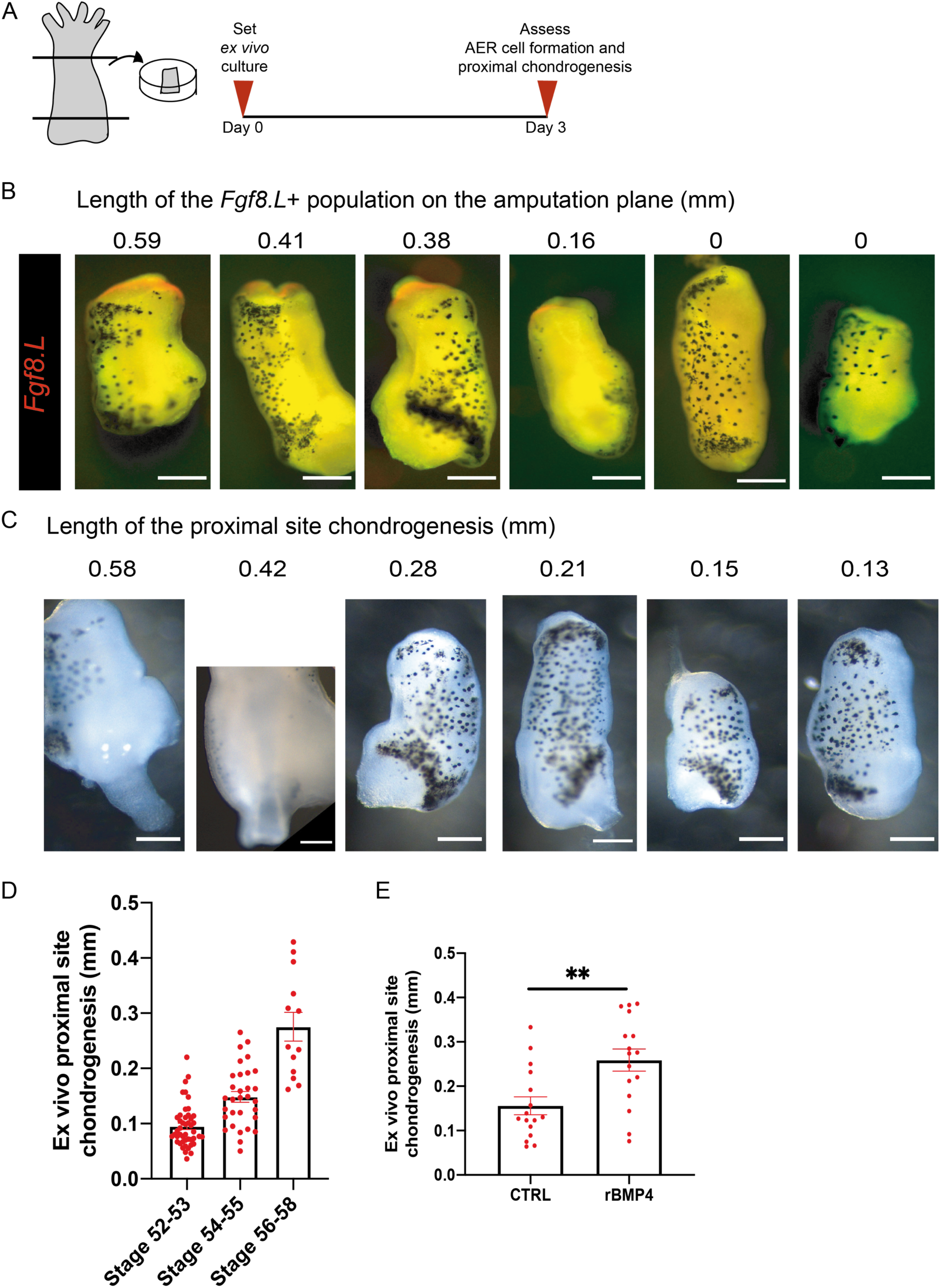
The distal site of *ex vivo* regenerating limbs can be used to detect AER cell formation, and the proximal site of explants can be used for detecting chondrogenesis. **(A)** Schematics describing the *ex vivo* culture protocol for assessing *Fgf8.L* mRNA expression at the distal site, and chondrogenesis levels at the proximal site. All assessments were carried out at 3-days post culture start. **(B)** Images of *Fgf8.L* stained limb explants at 3 dpa. Numbers at the top indicates AER cell formation measured as the length of *Fgf8.L*+ signal on the amputation plane. Red, *Fgf8.L* mRNA. Scale= 200 µm. (**C)** Images of chondrogenesis at the proximal site of explants at 3 dpa. Numbers at the top indicates the measured proximal chondrogenesis extent. Scale= 200 µm. **(D)** *Ex vivo* regenerating limb cultures can be used to investigate chondrogenesis. Explants were cultured for 3 days and chondrogenesis was measured as in (C). The extent of chondrogenesis seen at the proximal site of explants changes with the developmental stage and coincides with the progression of *in vivo* chondrogenesis (*5*). Regeneration-competent explants= total 46 samples from 4 biological replicates; Regeneration-restricted explants= total 31 samples from 3 biological replicates; Regeneration-incompetent explants= total 13 samples from 3 biological replicates. *P*\*\*< 0.001. **(E)** Explants were cultured for 3 days with BMP4 and the extent of chondrogenesis was measured. Addition of recombinant BMP4 to the explant media increased the observed chondrogenesis at the proximal site. Control 0.1% BSA, total n= 16 samples from 4 biological replicates; recombinant BMP4, total n= 16 samples from 4 biological replicates.

**Fig S13.**
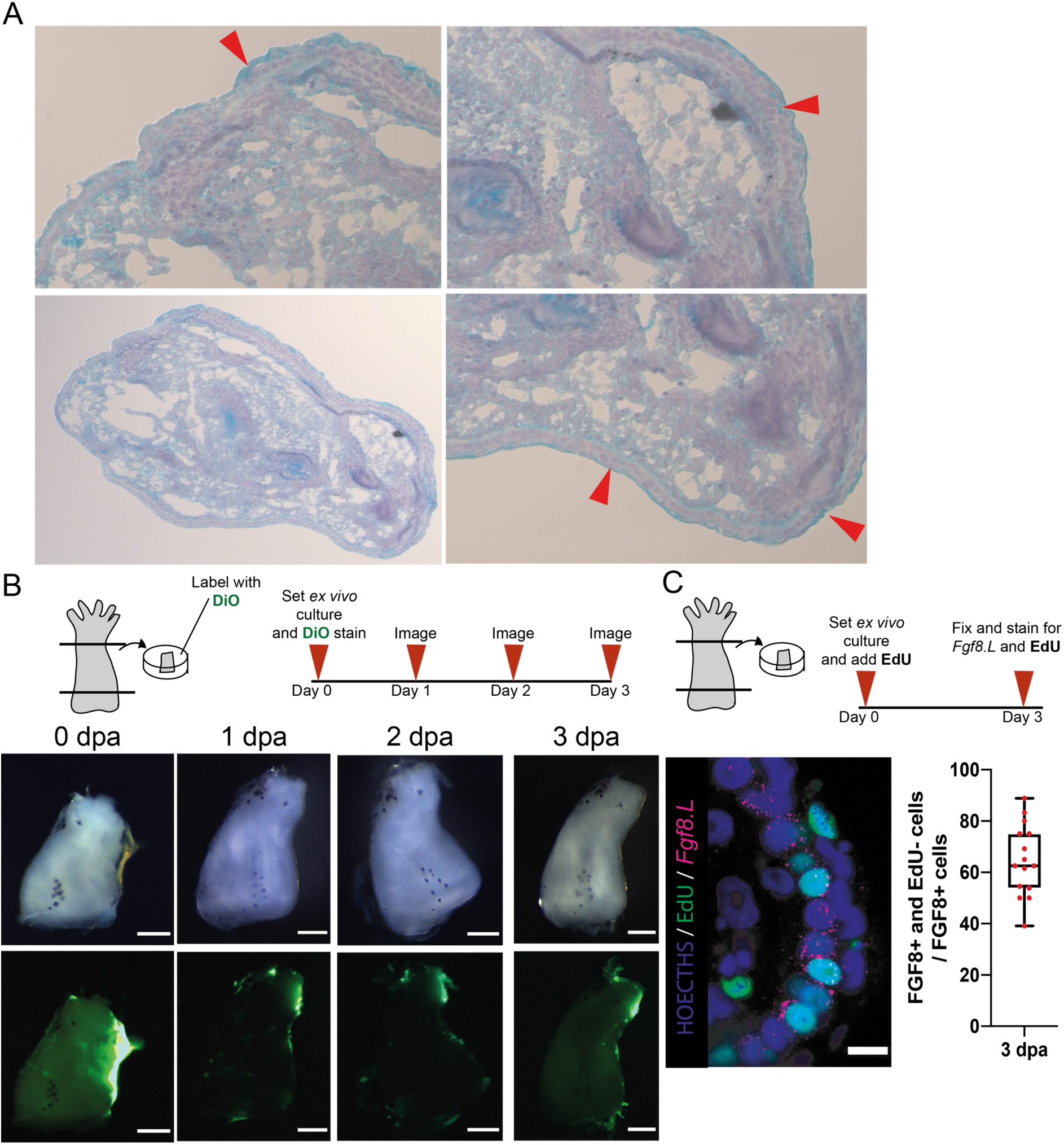
AER cells formation does not require cell division. **(A)** Explants are covered with cells morphologically similar to the surrounding basal epidermal cells as evidenced by haematoxylin, eosin, and Alcian blue stain. There are multi-layered or monolayered epidermal cells with cuboidal shape that can be seen not only at the distal site (right-bottom) but also at the lateral sides as well (right-top). A squamous layer can be seen above the basal epidermal cells. (**B)** (Top) Schematic describing DiO based tissue tracing. DiO labelling was performed after *ex vivo* cultures were harvested. Explants were imaged every day until day 3 in culture. (Bottom) DiO tracing applied to the sides of explants and traced over time and images were taken in brightfield and green channel. Traced tissues migrated to the distal and proximal amputation planes of explants. Total n = 22 from 2 biological replicates. Scale = 200 µm. **(C)** (Top) Schematics describing *ex vivo* culture with EdU treatment. EdU was added to explant media at the beginning of the culture. (Bottom-left) Explants were fixed and stained for *Fgf8.L* and EdU after day 3 in culture. (Bottom-middle) Example confocal image of a sample stained for *Fgf8.L, Lgr5.S*, and EdU, showing that not all *Fgf8.L+/Lgr5.S+* cells are EdU+. Hoechst, Blue; EdU, Green, *Fgf8.L* mRNA, Magenta. Scale = 10 µm. (Bottom-Right) Quantification of EdU positive AER cells proportion to all detected AER cells. 70% of AER cells are EdU negative. Total n= 15 from 3 biological replicates.

**Fig S14.**
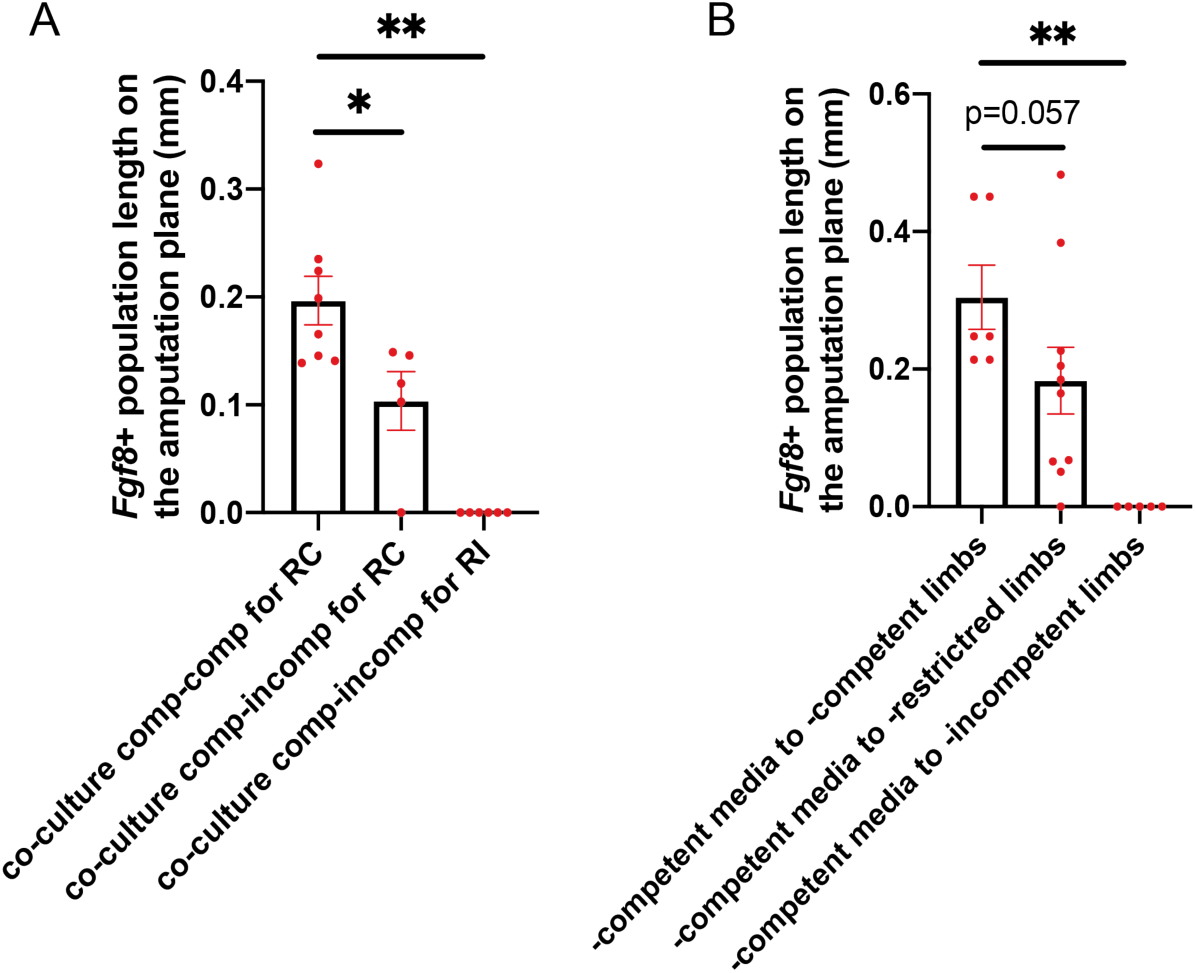
Regeneration-competent explants extrinsic cues are not sufficient to induce AER cells on the amputation plane of regeneration-incompetent explants. **(A)** Co-culturing regeneration-competent with-incompetent explants does not enable AER cell formation ability in -incompetent explants. Co-culture of regeneration-competent-competent and assess –competent: total n=8, from 2 biological replicates, Co-culture regeneration-competent-incompetent and assess –competent: n= 5 from 2 biological replicates. Co-culture regeneration-competent-incompetent for -incompetent total n= 6 from 2 biological replicates. *P*\*< 0.05, and *P*\*\*< 0.001. **(B)** Treatment with –competent conditioned media does not enable AER cell formation in -incompetent explants. Adding -competent-media to –competent explants: total n=6, from 2 biological replicates. Adding -competent-media to –restricted explants: n= 10, from 3 biological replicates. Adding -competent-media to –incompetent explants: n= 5, from 1 biological replicate. *P*\*\*< 0.001.

**Fig S15.**
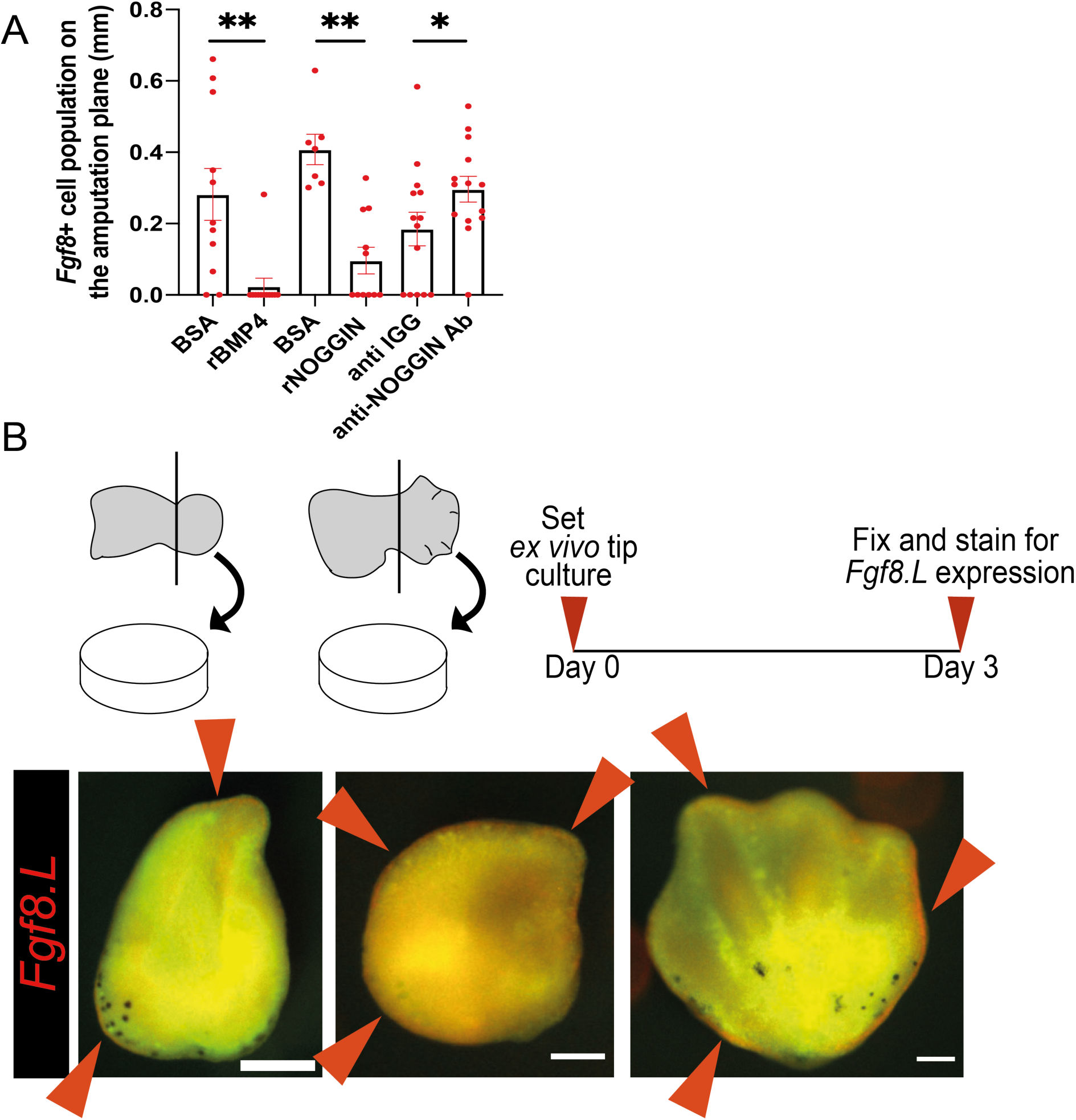
A regulated level of BMP pathway activation is required for AER cell formation, and reducing the proportion of chondrogenic lineage populations in explants can induce ectopic AER cell formation. **(A)** Regeneration-competent explants were treated with recombinant BMP4, recombinant NOGGIN, and anti-NOGGIN antibodies. Contralateral limbs were used as controls and treated with vehicle solutions (0.1% BSA, or anti-IGG). Recombinant BMP4 or NOGGIN additions block AER cell formation. Anti-NOGGIN antibody treatment enhances AER cell formation. From left to right 0.1% BSA: total n=11 from 3 biological replicates; rBMP4: total n= 12 from 3 biological replicates; 0.1% BSA: total n=7 from 2 biological replicates; rNOGGIN: total n=11 from 2 biological replicates; anti-IGG antibody: total n=14 from 4 biological replicates; anti-NOGGIN antibody: total n=14 from 4 biological replicates. Each sample group compared to their contralateral group to assess statistical significance. *P*\*< 0.05, and *P*\*\*< 0.001. **(B)** (Top) Schematic describing the protocol for culturing distal limb buds (NF stage ∼52) and early autopods (NF Stage ∼54). Tip explants were cultured for 3 days in explant media, and assessed for *Fgf8.L* expression. (Bottom) Tip cultures show ectopic *Fgf8.L* expression. Red arrows show *Fgf8.L* expression regions. Ectopic AER formation is seen in total 16/18 cases from 2 biological replicates. Red, *Fgf8.L* mRNA. Scale= 100 µm.

**Fig S16.**
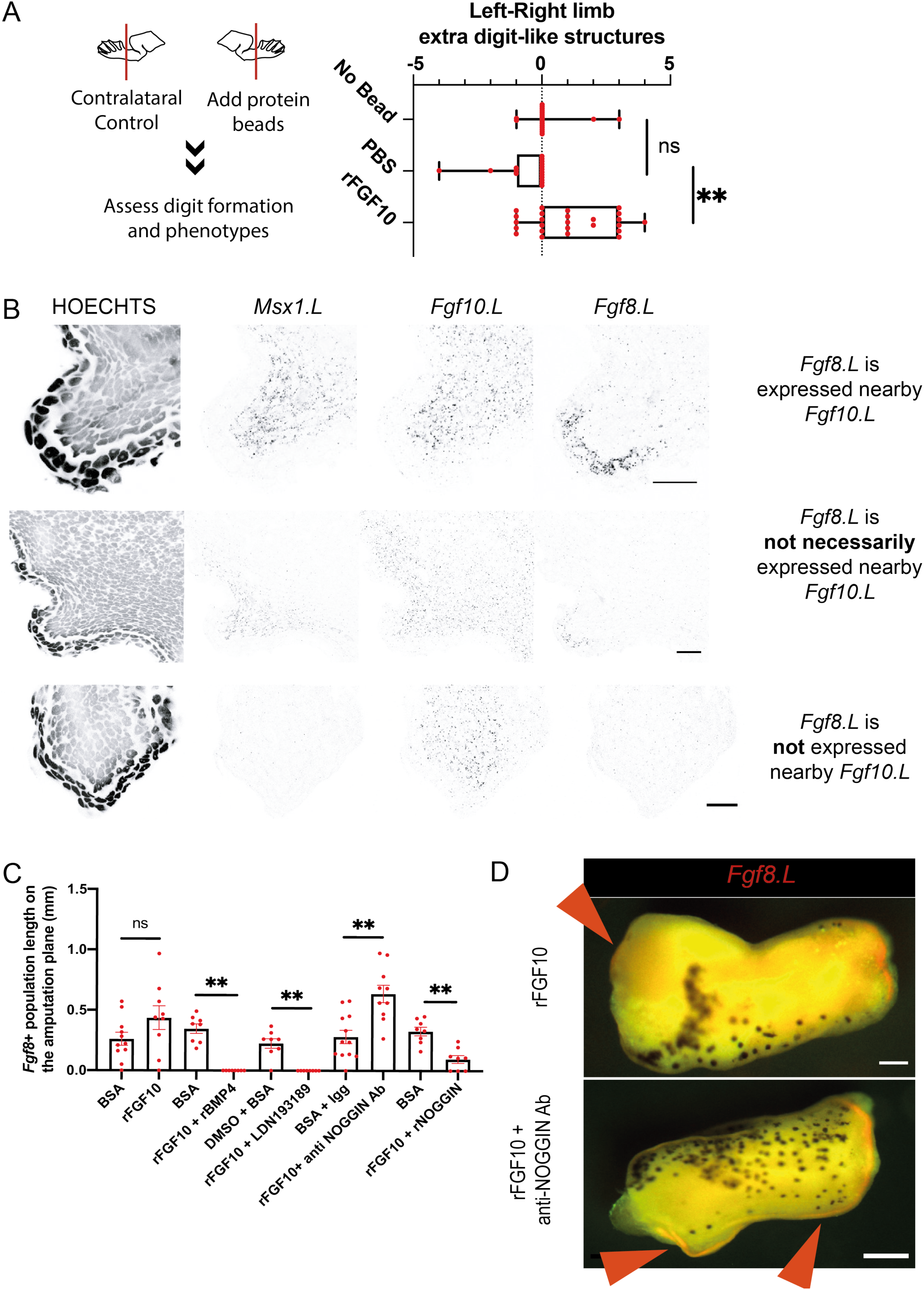
FGF10 application can restore regeneration and block chondrogenesis while FGF pathway inactivation enhances chondrogenesis. **(A)** Recombinant FGF10 application to distal amputations restore regeneration in –restricted and –incompetent tadpoles. –Restricted and –incompetent tadpole right and left hindlimbs were amputated and beads containing 0.1% BSA/PBS or recombinant FGF10 were placed on the right hindlimbs. Formed digits and digit-like structures were quantified in the right and left hindlimbs and the difference calculated. PBS beads application had no significant difference to empty controls, meanwhile recombinant FGF10 application improved regeneration. Empty total n=19 from 2 biological replicates; 0.1%/PBS bead total n=17 from 5 biological replicates; recombinant FGF10 bead total n=25 from 5 biological replicates. ns = not significant, *P*\*\*< 0.001. (**B)** Examples of confocal images of 5 dpa samples from regeneration-competent tadpoles stained for *Msx1.L, Fgf10.L* and *Fgf8.L*. Top image series show high levels of *Fgf10.L* and *Msx1.L* in the mesenchyme associated with high levels of *Fgf8.L* in the surrounding epidermis. Middle image series show that not all epidermis in proximity of *Fgf10.L* + mesenchymal cells are expressing *Fgf8.L. Msx1.L* + mesenchymal cells are more correlated to *Fgf8.L*+ epidermis than *Fgf10.L* + mesenchymal cells. Bottom, although there is a high level of *Fgf10* expression detected in mesenchyme, no *Fgf8.L* in epidermis or *Msx1.L* in mesenchyme can be seen. Scale, 20 µm. **(C)** Regeneration-competent explants were treated with rFGF10, alone or in combination with recombinant BMP4, recombinant NOGGIN, LDN193189, anti-NOGGIN antibody. 0.1% BSA/PBS and anti-IGG antibody were used as controls. From left to right, BSA: total n=11 from 2 biological replicates; FGF10: total n=9 from 2 biological replicates; BSA: total n=8 from 2 biological replicates; recombinant FGF10 and recombinant BMP4: total n=8 from 2 biological replicates; DMSO and BSA: total n=8 from 2 biological replicates; FGF10 and LDN total n=8 from 2 biological replicates; BSA and anti-IGG antibody: total n= 12 from 3 biological replicates; FGF10 and anti-NOGGIN antibody: total n=10 from 3 biological replicates; BSA: total n=8 from 2 biological replicates; recombinant FGF10 and recombinant NOGGIN: total n= 8 from 2 biological replicates. *P*\*< 0.05, and *P*\*\*< 0.001. **(D)** Example images of rFGF10 only or rFGF10 and anti-NOGGIN antibody treated explants showing ectopic *Fgf8.L* expression. rFgf10 treated explants can show a very mild expression of *Fgf8.L* at their proximal sites (n=4/7 from 2 biological replicates). rFGF10 and anti-NOGGIN antibody treated explants can show a substantial *Fgf8.L* expression at different sites of the explant (n=5/9 from 2 biological replicates). Scale, 200 µm.

**Fig S17.**
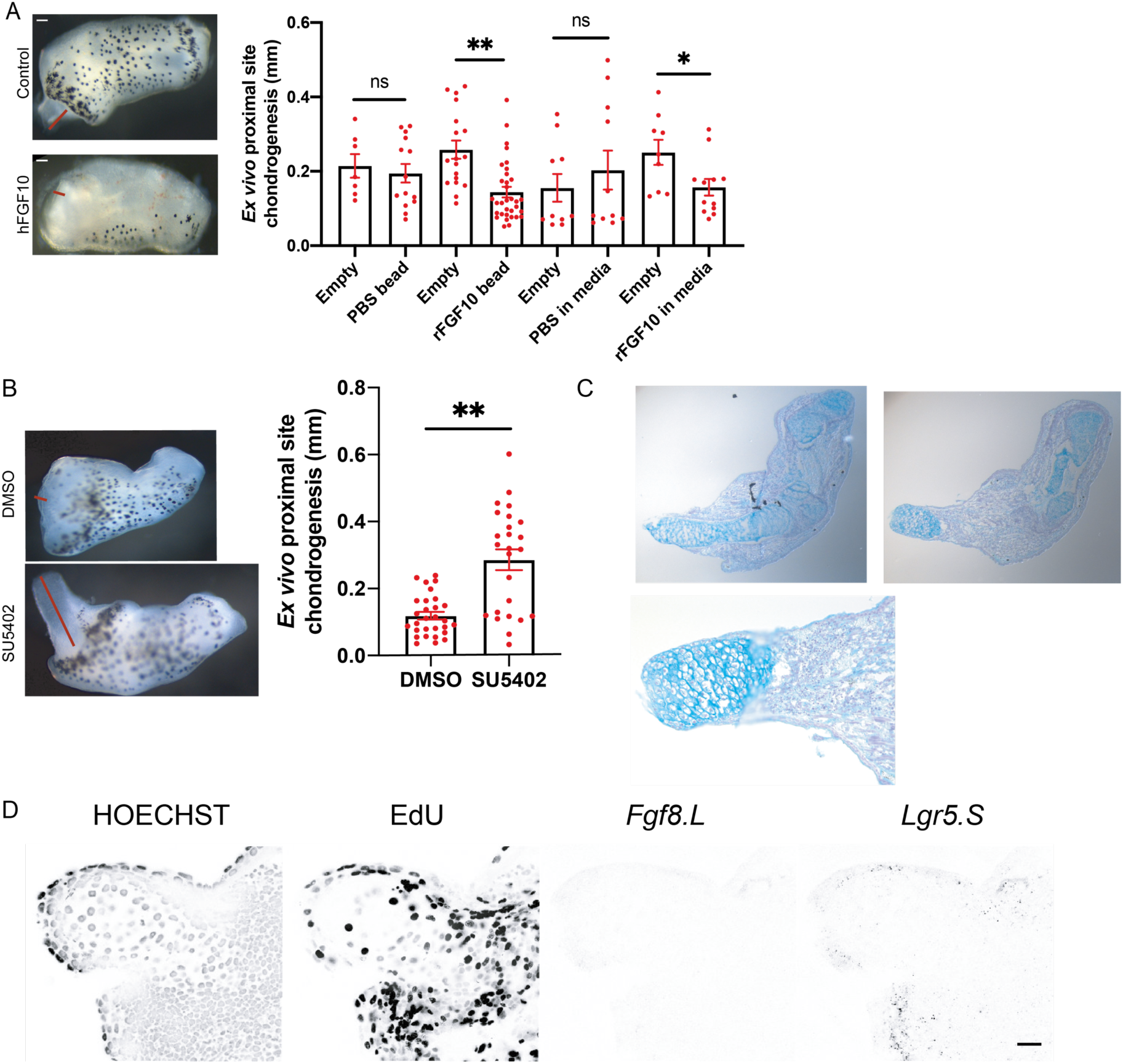
FGF10 application can block chondrogenesis, and blocking FGF pathway enhances chondrogenesis. **(A)** The effect of FGF10 on chondrogenesis is assessed by measuring the chondrogenic outgrowth at the proximal sites of -restricted explants at 3 dpa. Implanting 0.1% BSA/PBS beads to the proximal site or supplying 0.1% BSA/PBS to the media had no significant effect on chondrogenesis while implanting Fgf10 beads to the proximal site or supplying FGF10 in media reduced chondrogenesis. From left to right, empty and PBS beads total n ≥7, from at least 2 biological replicates; empty and FGF10 bead total n ≥ 14, from at least 4 biological replicates; empty and 0.1% BSA/PBS in media total n=10 from 3 biological replicates; empty and FGF10 in media ≥ n=14 from at least 3 biological replicates. ns = not significant, *P*\*< 0.05, and *P*\*\*< 0.001. **(B)** (Left) Example images of SU5402 treated explant showing extensive chondrogenesis at the proximal site. (Right) Blocking FGFR via small molecule inhibitor SU5402 extends chondrogenesis in 3 days for –competent and –restricted explants. Contralateral limbs were used as control and treated with DMSO. DMSO total n= 29, from 7 biological replicates, and SU5402 total n=25 from 7 biological replicates. *P*\*\*< 0.001. **(C)** Example histology images for explants treated with SU5402. The outgrowing structure are alcian blue rich indicative of chondrogenic cells. (**D)** Example confocal images of explants proximal site showing sparse circular nucleus indicative of chondrogenic cells as well as lack of *Fgf8.L* in epidermis. Scale bar = 15 µm.

**Supplemental table 1:** Quality control of scRNA-Seq data.

